# Unsupervised discovery of demixed, low-dimensional neural dynamics across multiple timescales through tensor components analysis

**DOI:** 10.1101/211128

**Authors:** Alex H. Williams, Tony Hyun Kim, Forea Wang, Saurabh Vyas, Stephen I. Ryu, Krishna V. Shenoy, Mark Schnitzer, Tamara G. Kolda, Surya Ganguli

## Abstract

Perceptions, thoughts and actions unfold over millisecond timescales, while learned behaviors can require many days to mature. While recent experimental advances enable large-scale and long-term neural recordings with high temporal fidelity, it remains a formidable challenge to extract unbiased and interpretable descriptions of how rapid single-trial circuit dynamics change slowly over many trials to mediate learning. We demonstrate a simple tensor components analysis (TCA) can meet this challenge by extracting three interconnected low dimensional descriptions of neural data: *neuron factors*, reflecting cell assemblies; *temporal factors*, reflecting rapid circuit dynamics mediating perceptions, thoughts, and actions within each trial; and *trial factors*, describing both long-term learning and trial-to-trial changes in cognitive state. We demonstrate the broad applicability of TCA by revealing insights into diverse datasets derived from artificial neural networks, large-scale calcium imaging of rodent prefrontal cortex during maze navigation, and multielectrode recordings of macaque motor cortex during brain machine interface learning.

## 1 Introduction

Two of the most challenging obstacles to understanding neural circuits are their diversity of dynamical timescales and the large number of neurons that contribute to their function. For instance, circuit dynamics mediating sensory perception, decision-making, attentional shifting, motor control, and higher cognition unfold over hundreds of milliseconds, while slower processes like motivation, long-term planning, and learning vary slowly, sometimes taking days or weeks to fully manifest [1–3]. Moreover, every execution of a behavior can involve the coordinated activity of extremely large neural populations, often distributed across multiple brain regions.

Recent experimental advances enable us to monitor all aspects of this biological complexity by recording large numbers of neurons [4–7] at high temporal precision [8] over long durations [9–11]. The resulting datasets can contain thousands of neural activity traces collected over thousands of behavioral trials. The genesis of such complex, large scale datasets now present a major data-analytic challenge to the field of neuroscience. Namely, how can we develop general purpose algorithms to extract from such complex data, simple and interpretable descriptions of collective circuit dynamics that underly not only rapid sensory, motor and cognitive acts, but also describe slower signatures of long-term planning and learning? Moreover, how can these algorithms operate in an unsupervised manner, to enable the discovery of novel and unexpected cognitive states that can vary on a trial by trial basis?

Neuroscientists have often turned to unbiased dimensionality reduction methods to understand these complex datasets [12, 13]. However, commonly used methods focus on reducing the complexity of fast, within-trial firing rate dynamics instead of extracting interpretable slow, across-trial structure. A common approach is to average neural activity across trials [13–15], thereby precluding the possibility of understanding of how cognition and behavior change on a trial by trial basis. More recent methods, including Gaussian Process Factor Analysis (GPFA) [16] and latent dynamical system models [17, 18], identify low-dimensional firing rate trajectories *within* each trial, but do not reduce the dimensionality *across* trials by extracting analogous low-dimensional trajectories over trials. Other works have separately focused on trial-to-trial variability in neural responses [19–22], and long-term trends across many trials [1, 3, 23–26], but without an explicit focus on obtaining simple low-dimensional descriptions. Thus, while current experimental data can simultaneously capture neural dynamics underlying both fast cognitive processes as well as slower learning processes, we lack general-purpose methods for extracting unbiased descriptions of both fast cognition and slower learning.

The most common and fundamental method for dimensionality reduction of neural data is Principal Components Analysis (PCA) [12, 13]. Here, we explore a simple extension of PCA that enables multi-timescale dimensionality reduction of neural dynamics both within trials and across trials. The key idea is to organize neural firing rates into a third-order tensor (i.e., a three-dimensional data table) with three axes corresponding to individual neurons (index 1), time within trial (index 2), and trial number (index 3). We then fit a tensor decomposition model (CANDECOMP/PARAFAC) [27, 28] to identify a set of low-dimensional components describing variability along each of these three axes. We refer to this procedure as Tensor Components Analysis (TCA).

We demonstrate that TCA yields insightful descriptions of a variety of neural datasets. In particular, it enables us to move beyond trial averaging by simultaneously identifying separate low-dimensional features for rapid within-trial neural dynamics and slower across-trial neural dynamics. Furthermore, as described below, TCA possesses a set of favorable theoretical properties that translate into significant interpretational advantages when applied to neural data. In particular, the components returned by TCA are often unique [29], unlike PCA which requires a biologically unrealistic orthogonality constraint to yield unique components. Because of the uniqueness of TCA, it achieves a demixing of neural data in which *individual* components are often in one-to-one correspondence with biologically interpretable variables. For example, as we see below, in diverse datasets, individual components correspond to sensations, decisions, actions, rewards and performance.

Below, after introducing the method, we show that TCA is equivalent to a form of multi-dimensional gain control and so can be interpreted as a generalization of a well-studied model of cortical function [30, 31]. We then give three examples of its utility. First, in an artificial neural circuit trained to solve the well-studied motion discrimination task [32], we show that TCA yields a simple one-dimensional description of the evolving connectivity and dynamics of the circuit during learning. Next, in a maze navigation task in rodents, we show that TCA can recover several aspects of trial structure and behavior, including perceptions, decisions, rewards, and errors, in an unsupervised, data-driven fashion. Finally, for a monkey operating a brain machine interface (BMI), we show that TCA extracts a simple view of motor learning when the BMI is altered to change the relationship between neural activity and motor action.

Thus, this work introduces a simple and broadly applicable method for identifying interpretable structure in multi-trial neural data, thereby providing a way to attack two of the most challenging aspects of modern large-scale neural recordings: their multi-timescale nature, and their high dimensionality. While TCA is a general-purpose method [33], we provide specialized code and step-by-step instructions for applying TCA to neural data, and describe how to interpret the outcomes of TCA within the context of systems neuroscience.

## 2 Results

### 2.1. Discovering multi-timescale structure through TCA

Before describing TCA, we first review the application of PCA for analyzing large-scale recordings. Consider a recording of *N* neurons over *K* experimental trials. We assume neural activity is recorded at *T* timepoints within each trial, but trials of variable duration can be aligned or temporally warped to accommodate this constraint (see, e.g., [34]). This dataset is naturally represented as an *N × T × K* array of firing rates, which is known in mathematics as a third-order tensor. Each element in this tensor, *x*_*ntk*_, denotes the firing rate of neuron *n* at time *t* within trial *k*. Here, the index *n* ranges from 1 to *N*, *t* ranges from 1 to *T*, and *k* ranges from 1 to *K*.

These datasets are very challenging to interpret in their raw format. Even nominally identical trials (e.g., neural responses elicited by repeats of an identical sensory stimulus) can exhibit significant trial-to-trial variability [22]. Under the assumption that such variability is simply irrelevant noise, a common method to simplify the table is to average across trials, obtaining a two dimensional table, or matrix, 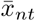, which holds the trial-averaged neural firing rates for every neuron *n* and timepoint *t* (fig. 1a). Even such a matrix can be difficult to understand in large-scale experiments containing many neurons and rich temporal dynamics. PCA summarizes these data by performing a decomposition into *R* components such that

**Fig 1.**
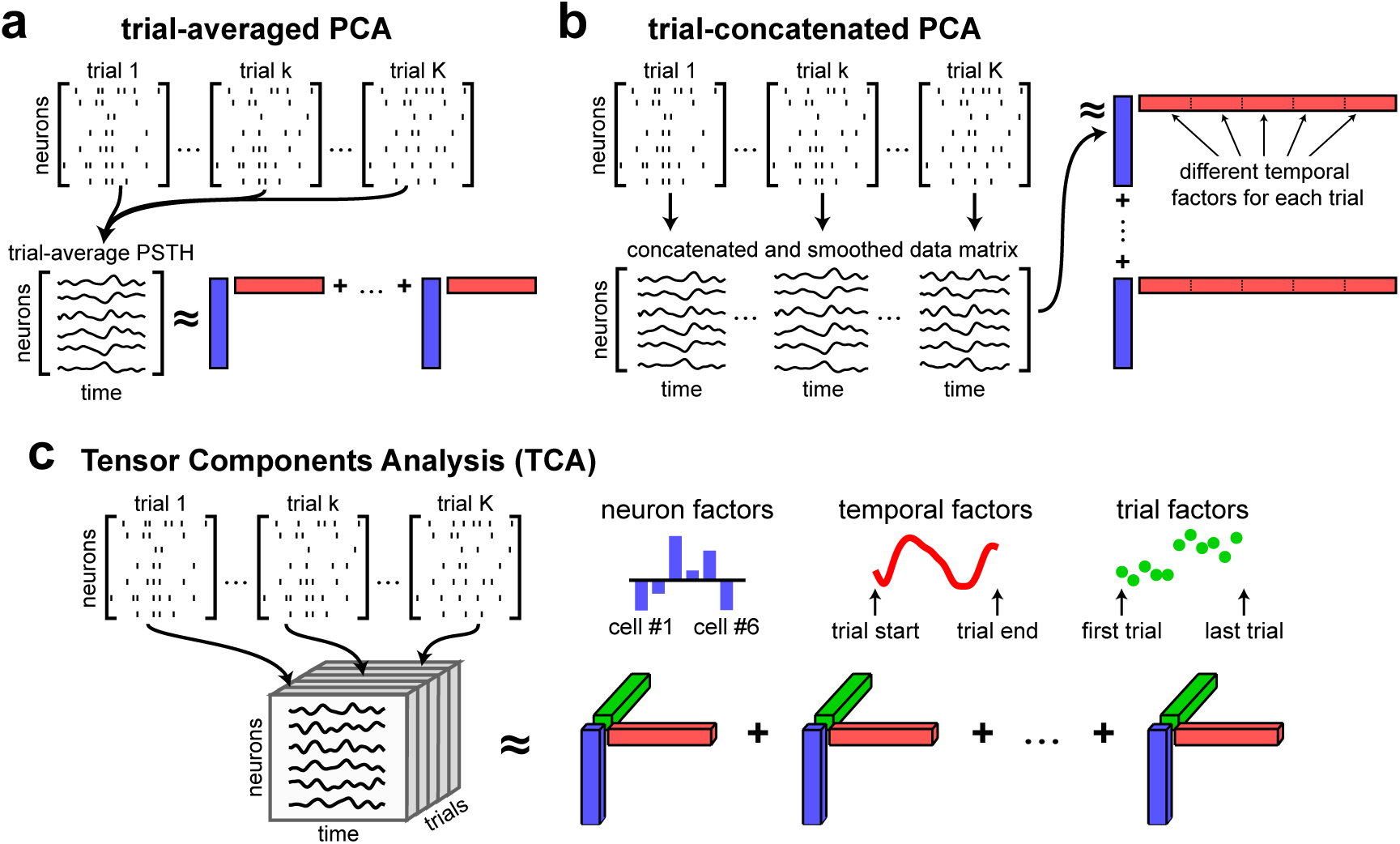
Tensor representation of trial-structured neural data. **(a)** Schematic of trial-averaged PCA for spiking data. The raw data is represented as a sequence of *N × T* matrices (top). These matrices are averaged across trials to build a matrix representation of neural firing rates. PCA approximates the trial-averaged matrix as a sum of outer products of vectors (see eq. (1)). Each outer product contains a neuron factor (blue rectangles) and a temporal factor (red rectangles). **(b)** Schematic of trial-concatenated PCA for spiking data. Raw data are temporally smoothed by a Gaussian filter to estimate neural firing rates before concatenating all trials along the time axis. Applying PCA produces a separate set of temporal factors for each trial (subsets of the red vectors). **(c)** Schematic of TCA. Raw data are smoothed and collected into a third order tensor with dimensions *N × T × K*. TCA approximates the data as a sum of outer products of three vectors, producing a third set of low-dimensional factors (trial factors, green vectors) that describe how activity changes across trials.

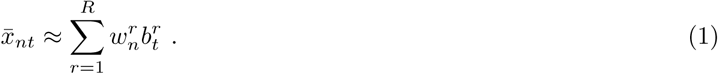

This decomposition projects the high-dimensional data (with *N* or *T* dimensions) into a low-dimensional space (with *R* dimensions). Each component, indexed by *r*, contains a coefficient across neurons, 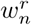, and a coefficient across timepoints, 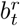. These terms can be collected into vectors: **w**^*r*^, of length *N*, which we call *neuron factors* (blue vectors in fig. 1), and **b**^*r*^, of length *T*, which we call *temporal factors* (red vectors in fig. 1). The neuron factors can be thought of as an ensemble of cells that exhibit correlated firing. The temporal factors can be thought of as a trial-averaged dynamical activity pattern for each ensemble. Overall, this *trial-averaged PCA* procedure reduces the original *N × T × K* datapoints into *R*(*N* + *T*) values, yielding a compact, and often insightful summary of the trial-averaged data [12, 13].

However, trial-averaging is motivated by the assumption that trial-to-trial variability is irrelevant noise, which is often at odds with our understanding of neural circuits and questions of experimental interest. For instance, even under repeated sensory stimuli, trial-to-trial variability may reflect fluctuations in interesting cognitive states, like attention or arousal [20, 21]. Also, under situations in which animals are learning a task, there will be systematic changes in neural dynamics over many trials, which would be rendered invisible by trial averaging. Intriguingly, as the field moves to study more complex tasks, we may find completely unexpected structured variability across trials, corresponding to different internal brain states on different trials. Ideally, we would like unbiased, data-driven methods to extract such dynamics simply by analyzing the data tensor.

One approach to retain the variability across trials is to concatenate multiple trials rather than averaging, thereby transforming the data tensor into an *N ×TK* matrix, and then applying PCA to this matrix (fig. 1b). This approach, which we call *trial-concatenated PCA*, is similar to Gaussian Process Factor Analysis (GPFA) [16], another specialized technique for neural data analysis. In trial-concatenated PCA, the *R* temporal factors are of length *TK* and do not enforce any commonality across trials. It therefore achieves a less significant reduction in the complexity of the data: the *NTK* numbers in the original data tensor are only reduced to *R*(*N* + *TK*) numbers, which can be cumbersome in experiments consisting of thousands of trials.

Our proposal is to directly deal with neural data in its natural third-order tensor format by performing a dimensionality reduction of this tensor (fig. 1c), rather than first converting it to a matrix. This tensor components analysis (TCA) method then yields the *R*-component decomposition [27, 28, 33]

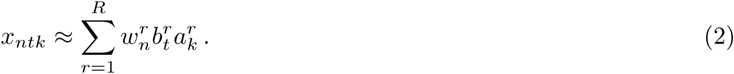

In analogy to PCA, we can think of **w**^*r*^ as a prototypical firing rate pattern across neurons, and we can think of **b**^*r*^ as a temporal basis function across time within trials. These neuron factors and temporal factors constitute structure that is common across all trials. We call the third set of factors, **a**^*r*^, *trial factors* (green vectors in fig. 1), which are new to TCA and not present in PCA. The trial factors can be thought of as trial-specific amplitudes for the within-trial activity patterns identified by the neuron and temporal factors. Thus, in TCA, the trial-to-trial fluctuations in neural activity are also embeded in *R*-dimensional space. TCA achieves a dramatic reduction of the original data tensor, reducing *NTK* datapoints to *R*(*N* + *T* + *K*) values, while still capturing trial-to-trial variability.

A subtle, but critical, difference between PCA and TCA is the uniqueness of the identified factors. In order to obtain unique factors, PCA constrains both the neuron and temporal factors to be orthogonal sets of vectors. This assumption is motivated by mathematical convenience rather than scientific principles. In real biological circuits, cell ensembles may overlap and temporal firing patterns may be correlated, producing non-orthogonal structure that is missed by PCA. In contrast, the TCA model often has a unique solution without further assumptions [29]. As we demonstrate below, TCA tends to extract non-orthogonal features of data that are more interpretable and meaningful than those extracted by PCA. In particular, we will see that TCA not only performs dimensionality reduction, but also demixing, by learning individual components that are in one-to-one correspondence with biologically interpretable variables.

### 2.2 TCA as a generalized cortical gain control model

Although TCA was originally developed as a statistical method [33], here we show that it concretely relates to a prominent theory of neural computation when applied to multi-trial datasets. In particular, performing TCA on neural data is equivalent to fitting a gain-modulated linear network model. In this network, *N* observed neurons (light gray circles, fig. 2a) are driven by a much smaller number of *R* unobserved, or latent, inputs (dark gray circles, fig. 2a) that have a fixed temporal profile but have varying amplitudes for each trial. The neuron factors of TCA, 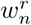 in eq. (2), correspond to the synaptic *weights* from each latent input *r* to each neuron *n*. The temporal factors of TCA, 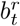, correspond to *basis functions* or the activity of input *r* at time *t*. Finally, the trial factors of TCA, 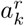 correspond to *amplitudes*, or gain, of latent input *r* on trial *k*. Such trial-to-trial fluctuations in amplitude have been observed in a variety of sensory systems [22, 35–37], and are believed to be an important and ubiquitous feature of cortical circuits [30, 31]. Furthermore, plausible cellular mechanisms for gain modulation have been examined by a number of experimental and computational studies [38–41]. The TCA model can be viewed as a higher, *R*-dimensional generalization of such theories. By allowing an *R*-dimensional space of possible gain modulations to different temporal factors, TCA can capture a rich diversity of changing multi-neuronal activity patterns across trials.

**Fig 2.**
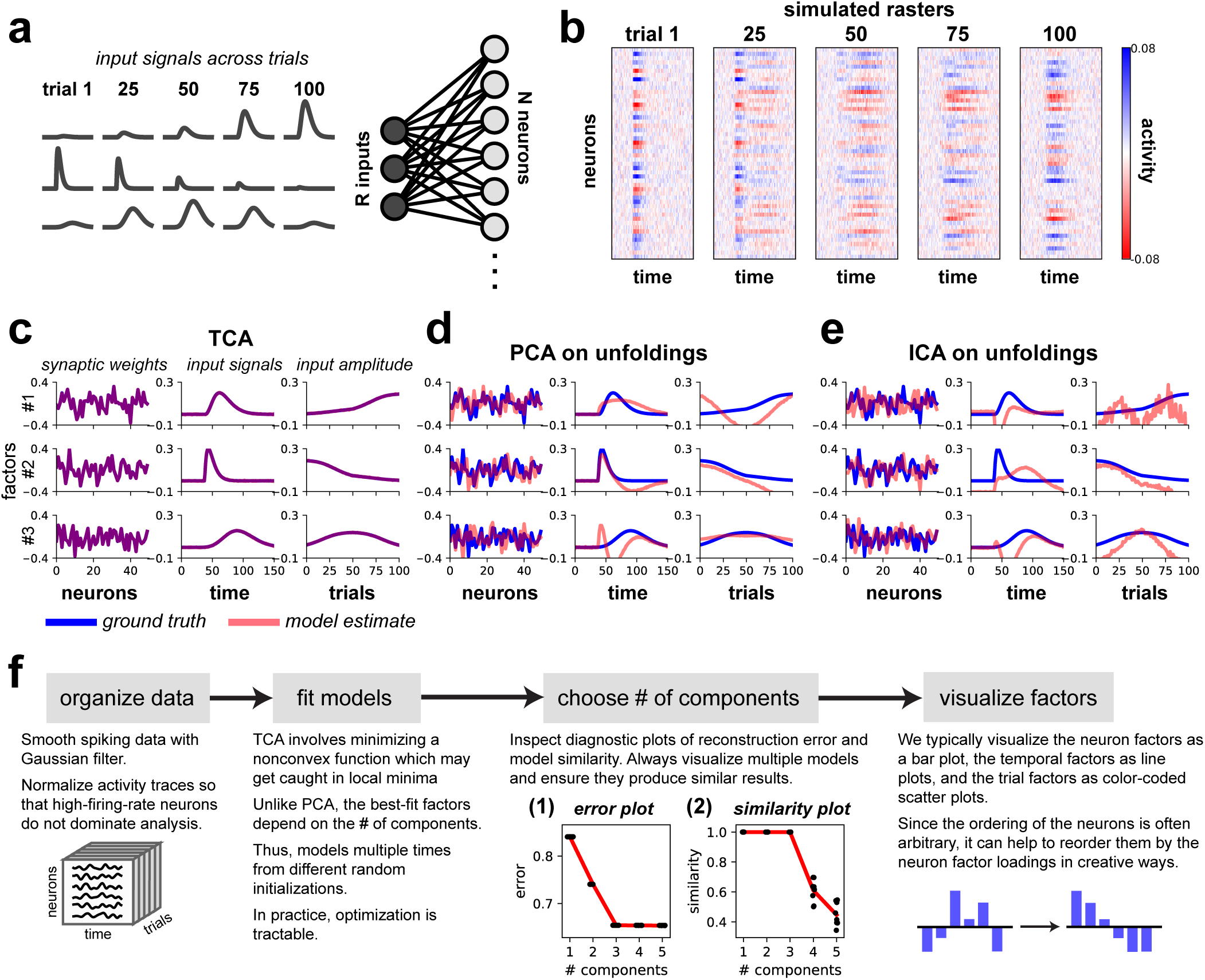
TCA precisely recovers the parameters of a linear model network. **(a)** Schematic of model network. Three input signals (dark gray) were delivered to a 1-layer, linear neural network with *N* = 50 neurons (light gray). Gaussian noise was added to the output units. **(b)** Simulated activity of all neurons on example trials. **(c)** The factors identified by 3-component TCA precisely match the network parameters. **(d-e)** Applying PCA (in **d**) or ICA (in **e**) to each of the tensor unfoldings does not recover the network parameters. **(f)** Analysis pipeline for TCA. **(f, inset 1)** Error plots showing normalized reconstruction error (vertical axis) for TCA models with different numbers of components (horizontal axis). The red line tracks the minimum error (i.e., best-fit model). Each black dot denotes a model fit from different initial parameters. All models fit from different initializations had essentially identical performance. Reconstruction error did not improve after more than 3 components were included. **(f, inset 2)** Similarity plot showing similarity score (eq. (12); vertical axis) for TCA models with different numbers of components (horizontal axis). Similarity for each model (black dot) is computed with respect to the best-fit model with the same number of components. The red line tracks the mean similarity as a function of the number of components. Adding more than 3 components caused models to be less reliably identified.

An important feature of TCA is that these network parameters can often be unambiguously identified from simulated data alone, due to the previously mentioned uniqueness property of TCA [29]. We confirmed this in a simple simulation with three latent inputs/components. In this example, the first component grows in amplitude across trials, the second component shrinks, and the third component grows and then shrinks in amplitude (fig. 2a). This model generates rich simulated population activity patterns across neurons, time, and trials as shown in (fig. 2b), where we have added Gaussian white noise to demonstrate the robustness of the method. When applied to noisy multi-neuronal traces, TCA with *R* = 3 components precisely extracted the network parameters (fig. 2c).

In contrast, neither PCA nor independent components analysis (ICA) [42] can recover the network parameters, as demonstrated in fig. 2d and fig. 2e respectively. Unlike TCA, both PCA and ICA are fundamentally matrix, not tensor, decomposition methods. Therefore they cannot be applied directly to the data tensor, but instead must be applied to three different matrices obtained by *tensor unfolding* (fig. 2 supp. 1; [33]). In essence, the unfolding procedure generalizes the trial-concatenated representation of the data tensor (fig. 1b) to allow concatenation across neurons or timepoints. This unfolding destroys natural structure across neurons, time, and trials in the third-order data tensor, thereby precluding the possibility of finding the ground truth synaptic weights, temporal basis functions, and trial amplitudes that actually generated observed neural activity patterns.

### 2.3 Choosing the number of components

A schematic view of the process of applying TCA to neural data is shown in fig. 2f (see *Methods* for more details). As in PCA and many other dimensionality reduction methods, a critical issue is the choice of the number of components, or dimensions *R*. We employ two methods to inform this choice. First, we inspect an *error plot* (fig. 2f, inset), which displays the model reconstruction error as a function of the number of components *R*. We normalize the reconstruction error to range between zero and one as described in section 4.5.1. This provides a metric analogous to the fraction of unexplained variance, which is used in PCA. As in PCA, a kink or leveling out in this plot indicates a point of diminishing returns for including more components. Unlike PCA, we run the optimization algorithm underlying TCA at each value of *R* multiple times from random initial conditions, and plot the normalized reconstruction error for all such models. Such repeated optimization runs enable us to check whether some runs converge to suboptimal solutions with high reconstruction error. As shown in (fig. 2f, inset), the error plot reveals that all runs at fixed *R* yield the same error, and moreover, the kink in the plot unambiguously reveals *R* = 3 as the true number of components in the generated data, in agreement with the ground truth.

A second method to assess the number of components involves generating a *similarity plot* (fig. 2f, inset), which displays how sensitive the recovered factors are to the initialization of the optimization procedure underlying TCA. For each component, we compute the similarity of all fitted models to the model with lowest reconstruction error by a similarity score bounded between zero (orthogonal factors) and one (identical factors). See section 4.5.1 for more details. Adding more components to the model can produce lower similarity scores, which complicates exploratory analysis since multiple low-dimensional descriptions may be consistent with the data. Like the error plot, the similarity plot unambiguously reveals *R* = 3 as the correct number of components, as decompositions with *R >* 3 are less consistent with each other (fig. 2f, inset). Notably, all models with *R* = 3 converge to *identical* components (up to permutations and re-scalings of factors), suggesting that only a single low-dimensional description, corresponding to the ground truth network parameters, achieves minimal reconstruction error. TCA consistently identifies this solution across multiple optimization runs.

### 2.4 TCA elucidates learning dynamics, circuit connectivity and computational mechanism in a nonlinear network

While TCA corresponds to a linear gain-modulated network, it can nevertheless reveal insights into the operation of more complex nonlinear networks, analogous to how PCA, a linear dimensionality reduction technique, allows visualization of low-dimensional nonlinear neural trajectories [12, 13]. We examine the application of TCA to nonlinear recurrent neural networks (RNNs), a powerful class of models that can learn to approximate any dynamical system [43]. RNNs have achieved success both in machine learning applications [44] and in modeling neural dynamics and behavior [45–48]. However, such models are so complex that they are often viewed as “black boxes.” Statistical methods that shed light on the function of RNNs and other complex computational models are therefore of great interest [49, 50]. Notably, while previous studies have focused on reverse-engineering RNNs with static parameters [51], few works have attempted to characterize how computational mechanisms in RNNs emerge over the process of learning, or optimization, of network parameters. Here we show TCA can naturally yield such a characterization for an RNN that learns to solve a simple sensory discrimination task, analogous to the well-known random dots direction-discrimination task [32].

Specifically, we trained an RNN with 50 neurons to estimate whether a noisy input signal had net positive or negative activity over a short time window, and indicate this estimate by exciting or inhibiting an output neuron (fig. 3a). We call trials with a net positive input *(+)-trials* and trials with a net negative input *(-)-trials*. The average amplitude of the input can be viewed as a proxy for the average motion energy of moving dots along a directional axis, with +/-corresponding to left/right, for example. The synaptic weights were updated by a simple gradient descent rule using backpropagation through time on a logistic loss function [52]. Within 750 trials the network performed the task with virtually 100% accuracy (fig. 3b).

**Fig 3.**
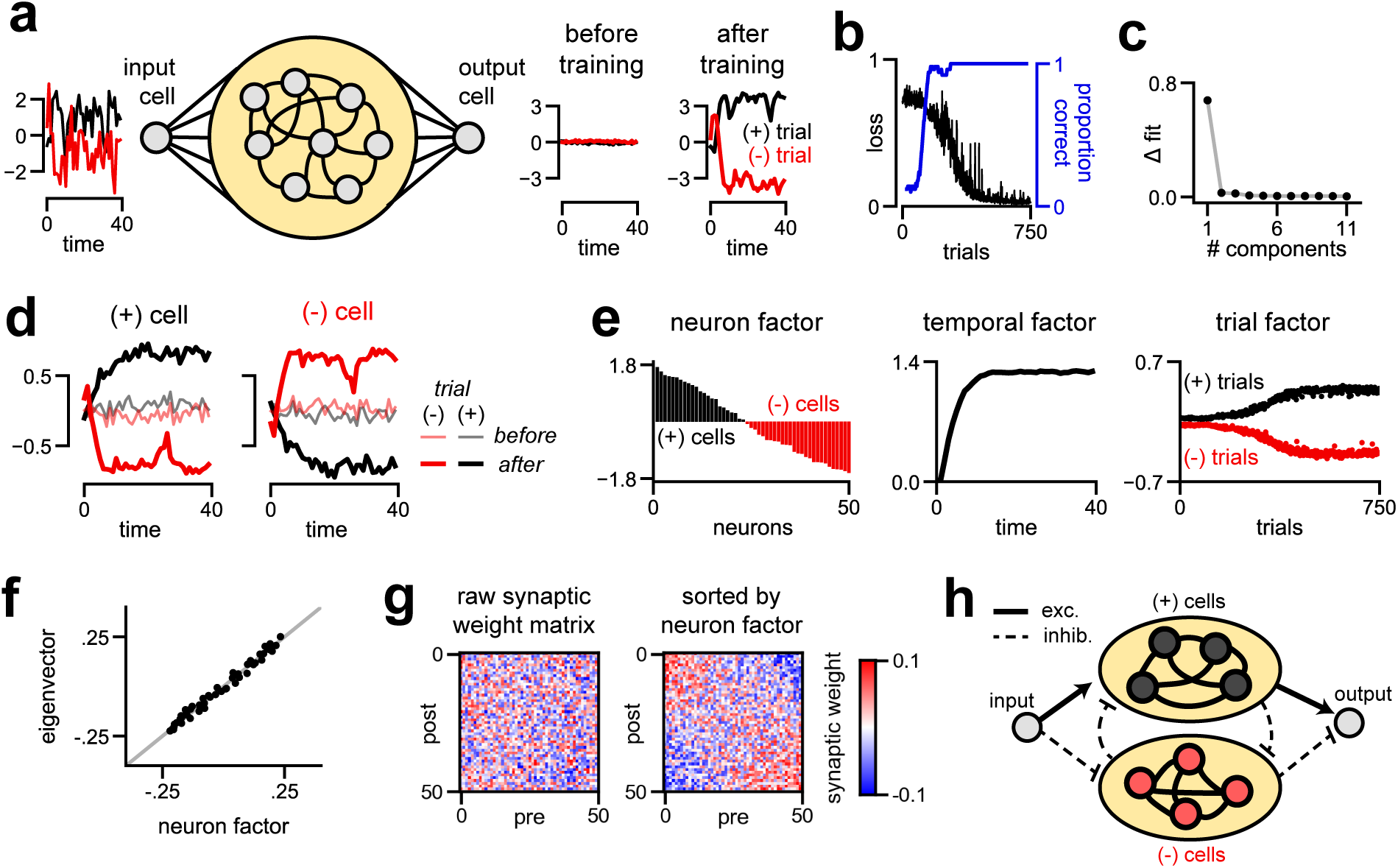
Unsupervised discovery of low-dimensional learning dynamics and mechanism in an model RNN. **(a)** Model schematic. A noisy input signal is broadcast to a recurrent population of neurons with all-to-all connectivity (yellow oval). On *(+)-trials* the input is net positive (black traces), while on *(-)-trials* the input is net negative (red traces). The network is trained to output the sign of the input signal with a large magnitude. **(b)** Learning curve for the model, showing the objective value on each trial over learning. **(c)** Scree plot showing the improvement in normalized reconstruction error as more components are added to the model. **(d)** An example *(+)-cell* and *(-)-cell* before and after training on both trial types. Black traces indicate *(+)-trials*, and red traces indicate *(-)-trials*. **(e)** Factors discovered by a one-component TCA applied simulated neuron activity over training. The neuron factor identifies *(+)-cells* (black bars) and *(-)-cells* (red bars), which have opposing correlations with the input signal. These two populations naturally exist in a randomly initialized network (trial 0), but become separated after during training, as described by the trial factor. **(f)** The neuron factor identified by TCA closely matches the principal eigenvector of the synaptic connectivity matrix post-learning. **(g)** The recurrent synaptic connectivity matrix post-learning. Resorting the neurons by their order in the neuron factor in **(e)** uncovers competitive connectivity between the *(+)-cells* and *(-)-cells*. **(h)** Simplified diagram of the learned mechanism for this network.

Remarkably, TCA needed only a single component to capture *both* the within-trial multi-neuronal circuit dynamics of decision making *and* the across-trial dynamics of learning. Adding more components led to negligible improvements in reconstruction error (fig. 3c). A single-component TCA model makes two strong predictions about this dataset. First, within all trials, the time course of evidence integration is shared across all neurons and is not substantially effected by training. Second, across trials, the amplitude of single cell responses are simply scaled by a common factor during learning. In essence, prior to learning, all cells have some small, random preference for one of the two input types, and learning corresponds to simply amplifying these initial tunings. We visually confirmed this prediction by examining single trial responses of individual cells. We observed two cell types within this model network: *(+)-cells* which were excited on (+)-trials and inhibited on (-)-trials (fig. 3d, left), and *(-)-cells* which were excited on (-)-trials and inhibited on (+)-trials (fig. 3d, right). The response amplitudes of both cell types magnified over learning, and typically the initial tuning (pale lines) aligned with the final tuning (dark lines). These trends are verified across the full population of cells in fig. 3 Supplement 1a-b.

We then visualized the three factors of the single-component TCA model (fig. 3e). We sorted the cells by their weight in the neuron factor, and plotted this factor, **w**^1^, as a bar plot (fig. 3e; left). Neurons with a positive weight are precisely the (+)-cells (black bars) defined earlier, while neurons with a negative weight were (-)-cells (red bars). While it is conceptually helpful to discretely categorize cells, the neuron factor illustrates that the model cells actually fall along a continuous spectrum rather than two discrete groups. The temporal basis function extracted by TCA, **b**^1^, reveals a common dynamical pattern within all trials corresponding to integration to a bound (fig. 3e; middle), similar to the example cells shown in Figure 3d. Finally, the trial factor of TCA, **a**^1^, recovered two important aspects of the neural dynamics (fig. 3e; right). First, the trial amplitude is positive for (+)-trials (black points) and negative for (-)-trials (red points), thereby providing a direct readout of the input on each trial. Second, over the course of learning, these two trial types become more separated, reflecting stronger internal responses to the stimulus and a more confident prediction at the output neuron. Intriguingly, this analysis reveals that the process of learning simply involves monotonically amplifying small but random initial selectivity for the +/-stimulus into a strong final selectivity.

This analysis also sheds light on the synaptic connectivity and computational mechanism of the RNN. To perform the task, the network must integrate evidence for the sign of the noisy stimulus over time. Linear model networks achieve this when the synaptic weight matrix has a single eigenvalue equal to one, and the remaining eigenvalues close to zero [53]. The eigenvector associated with this eigenvalue corresponds to a pattern of activity across neurons along which the network integrates evidence. The nonlinear RNN converged to a similar solution where one eigenvalue of the connectivity matrix is close to one, and the remaining eigenvalues are smaller and correspond to random noise in the synaptic connections (fig. 3, supp. 1a). Although the TCA model was fit only to the activity of the network, the prototypical firing pattern extracted by TCA in (fig. 3e; left) closely matched the principal eigenvector of the network’s synaptic connectivity matrix (fig. 3f). Thus, TCA extracted an important aspect of the network’s connectome from the raw simulated activity.

The neuron factor can also be used to better visualize and interpret the weight matrix itself. Since the original order of the neurons is arbitrary, the raw synaptic connectivity matrix appears to be unstructured noise (fig. 3g, left). However, re-sorting the neurons based on the neuron factor extracted by TCA, reveals a competitive connectivity between the (+)-cells and (-)-cells (fig. 3g, right). Specifically, neurons tend to send excitatory connections to cells in their same class, and inhibitory connections to cells of the opposite class. We also observed positive correlations between the neuron factor and the input and output synaptic weights of the network (fig. 3 supp. 1b-c). Taken together, these results provide a simple account of network function in which the input signal excites (+)-cells and inhibits (-)-cells on (+)-trials, and vice versa on (-)-trials. The two cell populations then compete for dominance in a winner-take-all fashion. Finally, the decision of the network is broadcast to the output cell by excitatory projections from the (+)-cells and inhibitory projections from the (-)-cells (fig. 3h).

In summary, TCA extracts a simple one-dimensional description of the activity of all neurons over all trials in this nonlinear network. Moreover, each of the three factors extracted by TCA have a simple neurobiological interpretation: the neuron factor **w**^1^ reveals a continuum of neurons interpolating between two cell assemblies, the temporal factor **b**^1^ describes the dominant neural activity underlying decision making, namely integration to a bound, and the trial amplitudes **a**^1^ reflect the trial-by-trial decisions of the network, as well as the long term amplification of stimulus selectivity underlying learning. Finally, even though the low-dimensional TCA factors were found in an unsupervised fashion from the raw neural activity, they provide direct insights into the synaptic connectivity and emergent computational mechanism underlying the network’s ability to learn and decide.

### 2.5 TCA compactly represents prefrontal activity during spatial navigation

Given the demonstrated success of TCA on an artificial nonlinear network, we next examined the performance of TCA on large-scale neurobiological datasets. We first examined the activity of cortical cells in mice performing a spatial navigation task with variable reward contingencies. A miniature microendoscope [54] was used to record fluorescence in GCaMP6m-expressing excitatory neurons in the medial prefrontal cortex while mice navigated a four-armed maze. Mice began each trial in either the east or west arm and chose to visit either the north or south arm, at which point a water reward was either dispensed or witheld (fig. 4a-b). We examined a dataset from a mouse containing *N* = 282 neurons recorded at *T* = 111 timepoints (at 10 Hz) on *K* = 600 behavioral trials, collected over a five day period. The rewarded navigational rules were switched periodically, prompting the mouse to explore different actions from each starting arm. Fluorescence traces for each neuron were shifted and scaled to range between zero and one in each session, and organized into a *N × T × K* tensor.

**Fig 4.**
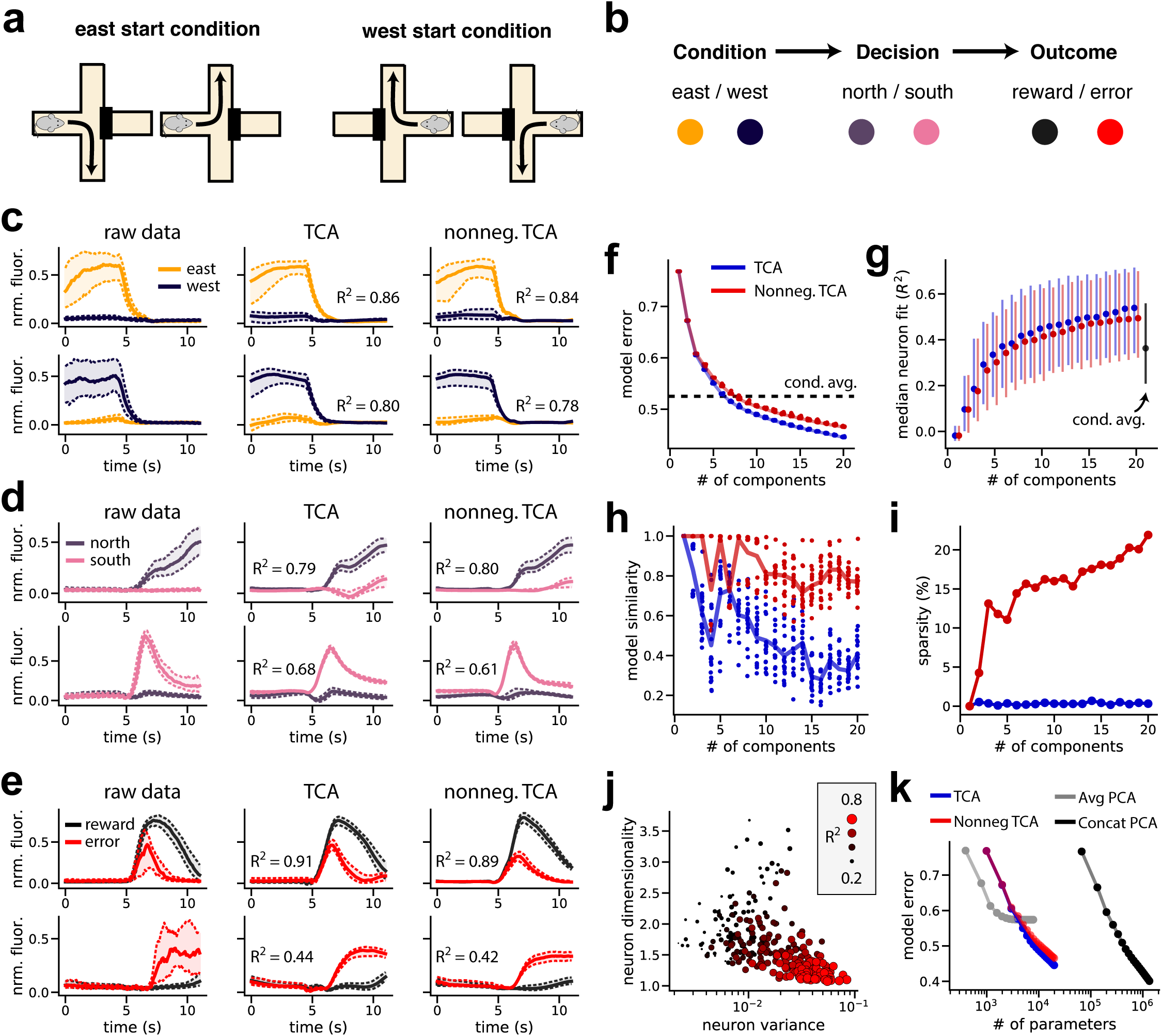
Reconstruction of single-cell activity during spatial navigation by unconstrained and nonnegative TCA. **(a)** All four possible combinations of starting and ending position on a trial. **(b)** Color scheme for three binary task variables (start location, end location, and reward). Each trial involves a sequential selection of these three variables. **(c)** Median fluorescence of example neurons that encode the starting location. Dashed lines denote upper and lower quartiles of the fluorescence. **(d-e)** Same as **(d)** but showing neurons that encode the ending location and the presence/absence of water reward. **(f)** Scree plot showing normalized reconstruction error for unconstrained (blue) and nonnegative (red) TCA, and the condition-averaged baseline model (black dashed line). Models were optimized from multiple initial parameters; each dot corresponds to a different optimization run. **(g)** Median coefficient of determination (*R*^2^) for neurons as a function of the number of model components for unconstrained TCA (blue), nonnegative TCA (red), and the condition-averaged baseline (black). Dots show the median *R*^2^ and the extent of the lines shows the first and third quartiles of the distribution. **(h)** Model similarity (section 4.5.1) as a function of model components for unconstrained (blue) and nonnegative (red) TCA. Each dot shows the similarity of a single optimization run compared to the best-fit model within each category. **(i)** Sparsity (proportion of zero elements) in the neuron factors of unconstrained (blue) and nonnegative decompositions. For each decomposition type, only the best-fit model is shown. **(j)** Neuron dimensionality (section 4.5.2) plotted against variance in activity. The size and color of the dots represent the *R*^2^ of a nonnegative decomposition with 15 components. **(k)** Normalized reconstruction error plotted against number of free parameters for trial-averaged PCA, trial-concatenated PCA, and TCA.

Neural firing in prefrontal cortical areas have previously been found to encode task variables, outcomes, value judgments, and cognitive strategies [25, 55–59]. We observed that many neurons selectively correlated with individual task variables on each trial: the initial arm of the maze (fig. 4c), the final arm (fig. 4d), and whether the mouse received a reward (fig. 4e). Notably, many of these neurons — particularly those with strong and robust coding properties — varied most strongly in amplitude across trials, suggesting that low-dimensional gain modulation is a reasonable model for these data. A TCA model with 15 components accurately modeled the activity of these individual cells and recovered their coding properties (fig. 4c-e; middle column; *R*^2^ between 0.44 and 0.91).

Since the fluorescence traces were normalized to be nonnegative, we also investigated the performance of *non-negative TCA*. This variant of TCA constrains the neuron, temporal, and trial factors to have nonnegative elements but is otherwise identical to standard TCA. Nonnegative TCA can produce more interpretable models, since the model is constrained to reconstruct the original data only through adding, but not subtracting, components, similar to nonnegative matrix factorization [60]. Despite this additional constraint, nonnegative TCA with 15 components reconstructed the activity of individual neurons with similar fidelity to an unconstrained TCA model (fig. 4c-e; right column; *R*^2^ between 0.42 and 0.89).

We then characterized the performance of TCA and nonnegative TCA across the full population of neurons. We compared both methods to a *condition-average baseline model*, which predicts the neural activity on each trial to be the trial-average population activity conditioned on the same trial trajectory (same starting arm and ending arm) and trial outcome (reward vs. error). That is, we computed the mean activity within each of the eight possible combinations of trial conditions, decisions, and outcomes, as used this to predict single-trial data. In essence, this baseline captures the average effect of all task variables, but does not account for trial-to-trial variability within each combination of task variables.

An error plot for unconstrained and nonnegative TCA showed three important findings (fig. 4f). First, nonnegative TCA had similar predictive performance to unconstrained TCA in terms of reconstruction error across all numbers of latent components (small gap between red and blue lines, fig. 4f). Second, both forms of TCA converged to very similar reconstruction error from twenty different random initializations, suggesting that the models did not get caught in highly suboptimal local minima during optimization (all blue points and all red points reached similar error, fig. 4f). Third, TCA models with more than 6 components matched or surpassed the condition-average baseline model, suggesting that relatively few components were needed to explain a substantial fraction of explainable variance in the dataset (dashed black line, fig. 4f). We also examined the performance of nonnegative and unconstrained TCA in terms of the *R*^2^ of individual neurons. Again, nonnegative TCA performed similarly to unconstrained TCA as judged by the median and upper/lower quartiles of the single neuron *R*^2^, and both models surpassed the simple condition-average baseline if they included more than 7 components (fig. 4g).

In addition to achieving similar accuracy to unconstrained TCA, nonnegative TCA possesses two important advantages. First, a similarity plot showed that nonnegative models converged more consistently to a similar set of low-dimensional components (fig. 4h). Second, the components recovered by nonnegative TCA were more sparse, meaning that each neuron’s activity across all trials was reconstructed by a smaller and more interpretable subset of components (fig. 4i).

While TCA could reconstruct the activity of many neurons very well (fig. 4c-e), other neurons were more difficult to fit (fig. 4, supp. 1). However, we observed that neurons with low *R*^2^ had firing patterns that were unreliably timed across trials and did not correlate with task variables (fig. 4, supp. 1b). To visualize this, we plotted the total variance and the dimensionality of each cell’s activity against the fit of a nonnegative TCA model with 15 components (fig. 4j). The dimensionality of each cell’s activity (see *Methods*, section 4.5.2) measures the trial-to-trial reliability of a cell’s firing: cells that fire consistently at the same time in each trial will be low-dimensional relative to cells that fire at different time points in each trial. First, this plot shows a negative correlation between variance and dimensionality: cells with higher variance (larger dynamic ranges in fluorescence) tended to be lower dimensional and thus more reliably timed across trials. Second, this plot shows these low-dimensional cells were well fit by TCA, suggesting that TCA summarizes the information encoded most reliably and strongly by this neural population. Moreover, outlier cells that defy a simple statistical characterization can be algorithmically identified and flagged for secondary analysis by sorting neurons by their *R*^2^ score under TCA.

TCA’s performance in summarizing neural population activity with very few parameters far exceeds that of trial-averaged PCA, which has sub-par performance, and trial-concatenated PCA, which requires many more parameters to achieve similar performance. This comparison is summarized in Figure 4k, which plots reconstruction error against the number of free parameters over 1 to 20 low-dimensional components for each class of models. Trial-averaged PCA (fig. 4k, gray line) has fewer parameters than TCA, but cannot account for trial-to-trial changes in activity, cannot achieve much lower than 60% error, and by construction entirely misses trial-to-trial fluctuations in neural firing that encode task variables. In contrast, trial-concatenated PCA (fig. 4k, black line) achieved comparable reconstruction error to TCA but required roughly 100x more free parameters, and is therefore much less interpretable. A TCA model with 15 components reduces the complexity of the data by 3 orders of magnitude, from ∼10^7^ datapoints to ∼10^4^ parameters; whereas a trial-concatenated PCA model with a comparable number of components only reduces the number of parameters to ∼10^6^.

### 2.6 Individual TCA components selectively correlate with individual task variables

These results demonstrate that TCA accurately describes the firing rates of single cells in a highly compact manner. We then examined whether this model identified an interpretable set of low-dimensional components. Figure 5 shows eight components from a 15-component nonnegative TCA model (the remaining seven factors carry similar information and are shown in fig. 5, supp. 1). Each nonnegative TCA component identified a sub-population, or assembly of cells (neuron factor; left column) with a common intra-trial temporal dynamics (temporal factor; middle column) that was differentially activated across trials (trial factor; right column).

**Fig 5.**
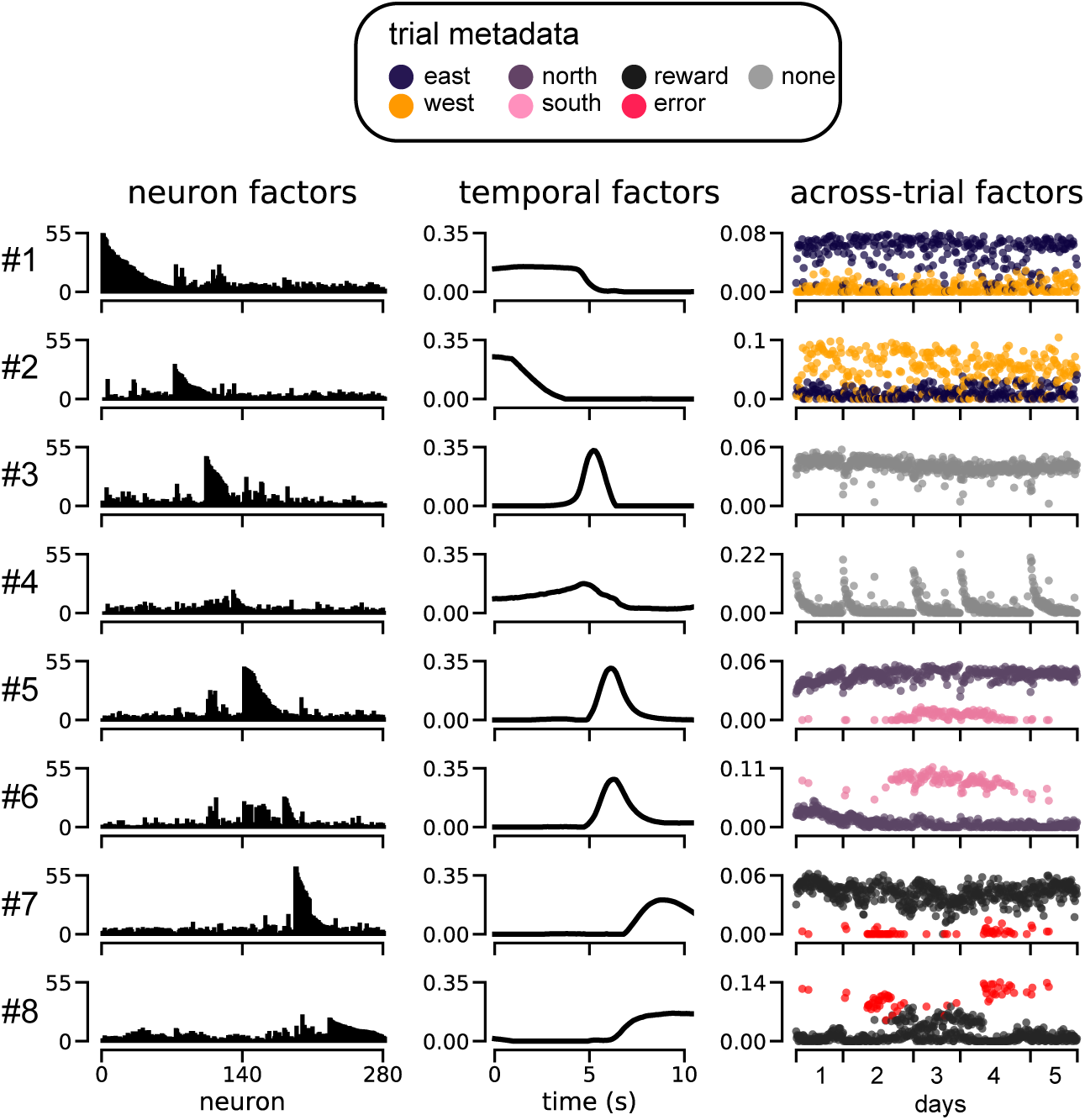
Nonnegative TCA of prefrontal cortical activity during spatial navigation. Eight low-dimensional components, each containing a neuron factor (left column), temporal factor (middle column), and trial factor (right column) are shown from a 15-component model (see fig. 5, supp. 1 for the remaining seven components). For each component, the trial factor is color-coded by the task variable it is most highly correlated with.

In contrast, PCA identified factors that contained complex mixtures of coding for the mouse’s position, choice, and reward on each trial (fig. 5, supp. 2), hampering interpretability [34]. TCA on the other hand, isolated each of these task variables into separate components: each trial factor selectively correlated with a *single* task variable, as indicated by the color-coded scatterplots in fig. 5. Overall, the TCA model uncovers, in a completely unsupervised manner, a compelling qualitative view of prefrontal dynamics in which largely distinct subsets of neurons (fig. 5, left columns) are active at successive times within a trial (fig. 5 middle column) and whose variation across trials (fig. 5 right column) encodes a highly interpretable single task variable.

Specifically, components 1-2 uncover neurons that encode the starting location (component 1, east trials; component 2, west trials), components 5-6 encode the destination arm (component 5, north trials; component 6, south trials), and components 7-8 encode the trial outcome (component 7, rewarded trials; component 8, error trials). Interestingly, the temporal factors indicate that these components are sequentially activated in each trial: components 1-2 activated before components 5-6, which in turn activated before components 7-8, in agreement with the schematic flow diagram shown in fig. 4b.

Intriguingly, TCA also uncovers unexpected components, like components 3-4 which activate prior to the destination and outcome-related components (i.e., components 5-8). Component 4 displays systematic reductions in activity across trials within each day, while component 3 is active on nearly every single trial. Component 4 could potentially correspond, for example, to a novelty or arousal signal that wanes over trials within a day. While further experiments will be required to ascertain whether this interpretation is correct, the extraction of these components illustrates the potential power of TCA as an unbiased exploratory data analysis technique to extract unobserved cognitive states and separate them from observable aspects of trial-to-trial variations in behavior.

It is important to emphasize that TCA is an unsupervised method that only has access to the neural data tensor, and does not receive any information about task variables like starting location, ending location, and reward. Therefore, the correspondence between TCA trial factors and behavioral information demonstrated in fig. 5, constitutes an unbiased revelation of task structure directly from neural data. Moreover, individual components extracted by TCA are in one-to-one correspondence with meaningful aspects of task structure and behavior, a property not shared by many other dimensionality reduction algorithms.

### 2.7 TCA reveals two-dimensional learning dynamics in macaque motor cortex after a BMI perturbation

In the previous section we validated TCA on a dataset where the animal’s behavior decomposed into a set of discrete experimental conditions, choices, and trial outcomes. We next applied this method to a brain-machine interface (BMI) learning task, in which the behavior on each trial was quantified by a continuous path of a computer cursor. The cursor movement on each trial is never identical and is difficult to summarize concisely in a principled manner. In these more unstructured scenarios, supervised methods such as classification and regression can be difficult to construct, making unsupervised dimensionality reduction methods an important tool to explore hypotheses and let the low-dimensional structure of the data “speak for itself.”

Specifically, we collected multi-unit data from the pre-motor and primary motor cortices of a Rhesus macaque (*Macaca mulatta*) controlling a computer cursor in a 2D plane through a brain-machine interface (fig. 6a). Spikes were recorded when the voltage signal crossed below −4.5 times the root-mean-square voltage. The monkey was trained to make point-to-point reaches from a central position to one of eight radial targets. For simplicity, we initially investigated neural activity during 45° outward reaches. The cursor velocity was controlled by a velocity Kalman filter decoder, which was driven by non-sorted multi-unit activity (-4.5 root-mean-square threshold crossings) and fit using relations between neural activity and reaches by the monkey’s contralateral arm at the beginning of the experiment [61]. We analyzed multi-unit activity during subsequent reaches, which used this decoder as a BMI interface directly from neural activity to cursor motion. These initial reaches were accurate (fig. 6b, left) and took less than one second to execute (fig. 6c, first 30 trials).

**Fig 6.**
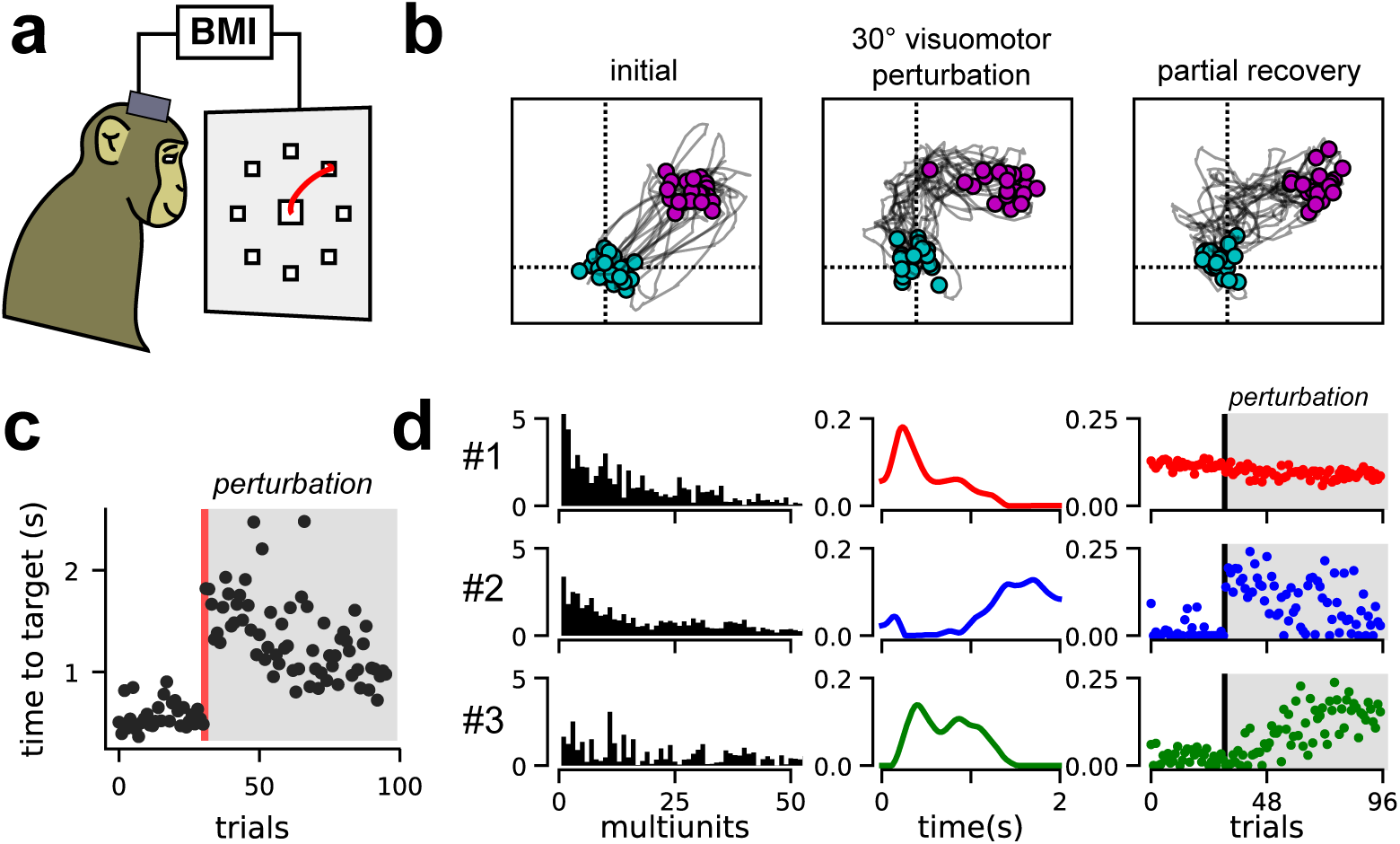
TCA reveals two-dimensional learning dynamics in primate motor cortex during BMI cursor control. **(a)** Schematic of monkey making center-out, point-to-point reaches in BMI task. **(b)** Cursor trajectories to a 45° target position. Twenty trials are shown at three stages of the behavioral session showing initial performance (left), performance immediately after a 30° counterclockwise visuomotor perturbation (middle), and performance after learning, at the end of the behavioral session. Cyan and magenta points respectively denote the cursor position at the beginning and end of the trial. **(c)** Time for the cursor to reach target for each trial in seconds. The visuomotor perturbation was introduced after 31 trials (red line). **(d)** An optimal 3-component nonnegative TCA on smoothed multi-unit spike trains recorded from motor cortex during virtual reaches reveals two components (2-3) that capture learning after the BMI perturbation.

We then perturbed the BMI decoder by rotating the output cursor velocities counterclockwise by 30° (a visuomotor rotation). Thus, the same neural activity pattern that originally caused a motion of the cursor towards the 45° direction, now caused a maladaptive motion in the 75° direction, yielding an immediate drop in performance: the cursor trajectories were biased in the counterclockwise direction (fig. 6b, middle), and took longer to reach the target (fig. 6c, trials following perturbation). These deficits were partially recovered within a single training session as the monkey adapted to the new decoder. By the end of the session, the monkey made more direct cursor movements (fig. 6b, right) and achieved the target more quickly (fig. 6c).

We applied TCA and nonnegative TCA to the raw spike trains smoothed with a Gaussian filter with a standard devation of 50 ms [34]. We again found that nonnegative TCA fit the data with similar reconstruction error and higher reliability than unconstrained TCA (fig. 6 supp. 1). To examine a simple account of learning dynamics, we examined a nonnegative TCA model with 3 components. Models with fewer than 3 components had substantially worse reconstruction error, while models with more components had only moderately better performance and occasionally converged to dissimilar parameters during optimization (fig. 6 supp. 1).

The neuron, temporal, and trial factors of the nonnegative TCA model are shown in Figure 6d. Component 1 (red) described multi-units that were active at the beginning of each trial, and were consistently active over all trials. The other two components described multi-units that were inactive before the BMI perturbation, and became active only after the perturbation, thereby capturing motor learning. Component 2 (blue) became active on trials immediately after the BMI perturbation, but then slowly decayed over successive trials. Within a single trial, this component was only active at late stages in the reach. Component 3 (green) on the other-hand was not active on trials immediately following the BMI perturbation, but did activate slowly across successive trials. Within a single trial, this component was active earlier in the reach. These results suggest a mode of motor learning in which a suboptimal, late reaching-stage correction is initially used to perform the task (component 2). Over time, this component is slowly traded for a more optimal early reaching-stage correction (component 3). Interestingly, motor learning did not involve extinguishing neural dynamics present before the perturbation (component 1), even though this component is maladaptive after the perturbation.

We were able to confirm this intuition by relating each of these components to a different phase of motor execution and learning. Figure 7a plots cursor trajectories on individual reaches before the perturbation (left), immediately following the perturbation (middle), and at the end of the behavioral session (right). Every 50ms the trajectory was colored based on the component with the largest activation at that timepoint and trial. Prior to the perturbation, component 1 (red) dominated; the other two components were nearly inactive since their TCA trial factor amplitudes were near zero before the perturbation (see fig. 6d). Immediately following the perturbation, component 1 still dominated in the early phase of each trial, producing a counterclockwise off-target trajectory. However, component 2 dominated the second half of each trial at which point the monkey performed a “corrective” horizontal movement to compensate for the initial error. Finally, near the end of the training session, component 3 was most active at many stages of the reach. Typically, the cursor moved directly towards the 45° target when component 3 was active, suggesting that component 3 captured learned neural dynamics that were correctly adapted to the perturbed visuomotor environment.

**Fig 7.**
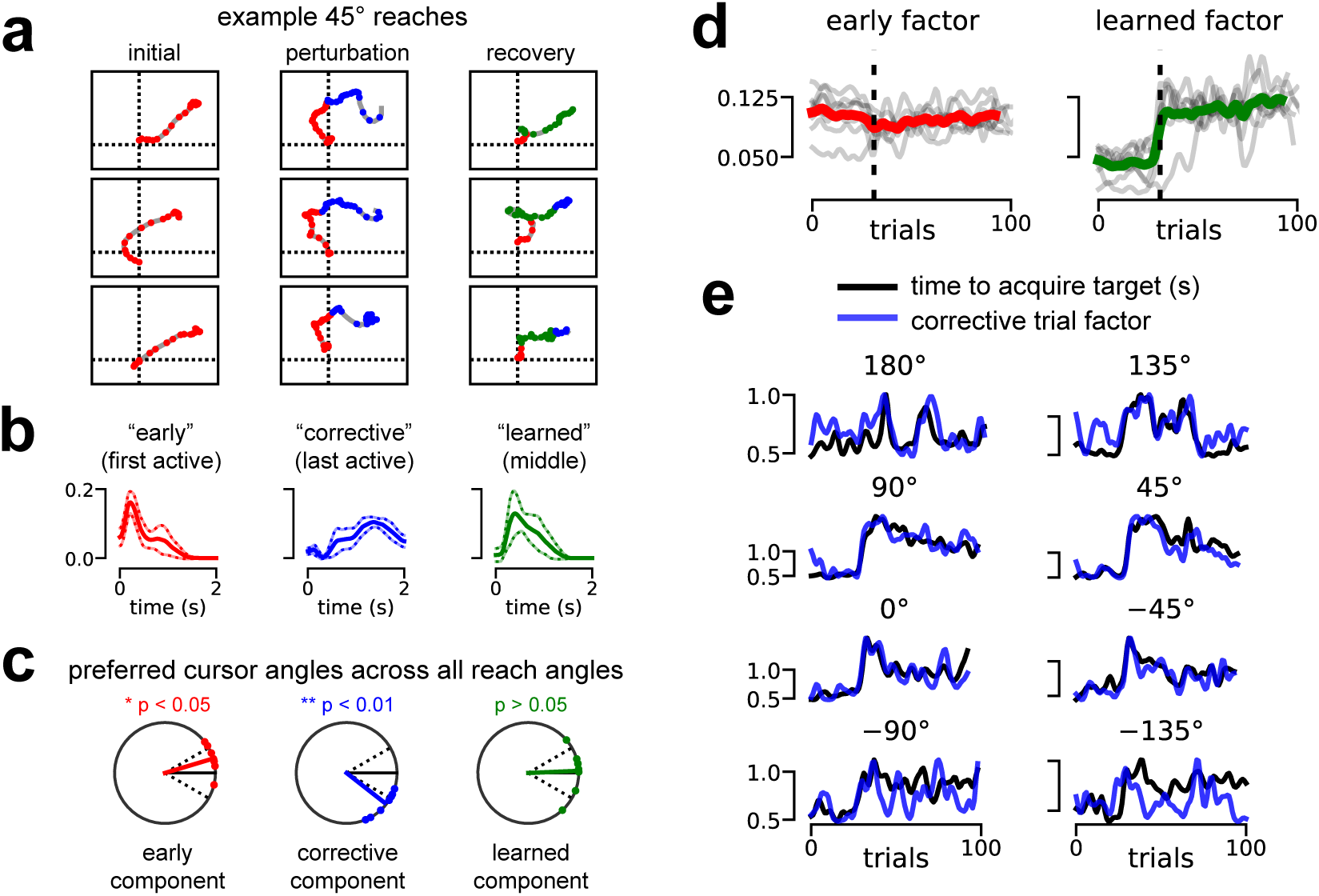
TCA tracks performance and uncovers “corrective” dynamics in BMI adaptation task. **(a)** Cursor trajectories for 45° cursor reaches. Every 50 ms, the trajectory is colored by the TCA component with the strongest activation at that timepoint and trial. Components were colored according to the definition in panel (b). Three example trajectories are shown at three stages of the experiment: reaches before the visuomotor perturbation (left), reaches immediately following the perturbation (middle), and reaches at the end of the behavioral session. **(b)** Average low-dimensional temporal factors identified by nonnegative TCA across all eight reach angles. The *early component* had the earliest active temporal factor (red). The *corrective component* had the last active temporal factor (blue). The *learned component* was the second active temporal factor (green). Solid and dashed lines denote mean +/- standard deviation. **(c)** Preferred cursor angles for each component type after the visuomotor perturbation. All data were rotated so that the target reach angle was at 0° (solid black line). Dashed black lines denote +/- 30° for reference, which was the magnitude of the visuomotor perturbation. On average, the *early component* was associated with a cursor angle misaligned counterclockwise from the target (red). The *corrective component* preferred angle was aligned clockwise from the target (blue) by about 30°, in a way that could compensate for the 30° counterclockwise misalignment of the *early component*. The *learned component* preferred angle not significantly different from that of the actual target. **(d)** Smoothed trial factors for the *early component* and *learned component*. Colored lines denote averages across all reach angles; gray lines denote the factors for each of the eight reach conditions. Factors were smoothed with a Gaussian filter with 1.5 standard deviation for visualization purposes. **(e)** Smoothed trial factor for the *corrective component* (blue) and smoothed behavioral performance (black) quantified by seconds to reach target. Each subplot shows data for a different reach angle. All signals were smoothed with a Gaussian filter with 1.5 standard deviation for visualization.

Based on these observations, we called the component active at the beginning of each trial the *early component* (#1 in fig. 6), the component active at the end of each trial the *corrective component* (#2 in fig. 6), and the component active in the middle of each trial the *learned component* (#3 in fig. 6). These components are colored red, blue, and green respectively in both Figure 6 and Figure 7. We then fit 3-component TCA models separately to each of the eight reach angles, and operationally defined the components as *early*, *corrective*, and *learned* based on the peak magnitude of their associated within-trial temporal basis functions (fig. 7b). This very simple definition yielded similar interpretations for low-dimensional components separately fit across different reach angles.

Similar to computing a directional tuning curve for an individual neuron [62], we examined the *preferred cursor angles* of each low-dimensional component by computing the average cursor velocity weighted by activity of the component (see *Methods*, section 4.4.7). To compare across all target reach angles, we rotated the preferred angles so that the target was situated at 0° (black line, fig. 7c). All preferred angles were computed on post-perturbation trials. When the *early component* was active, the cursor typically moved at an angle counterclockwise to the target (*p <* 0.05, one sample test for the mean angle), reflecting our previous observation that the early component encodes pre-perturbation dynamics that are maladaptive post-perturbation (fig. 7c, left). When the *corrective component* was active, the cursor typically moved at an angle clockwise to the target (*p <* 0.01, one sample test for the mean angle), reflecting a late-trial compensation for the error introduced by the early component (fig. 7c, middle). Finally, the *learned component* was not significantly different from the target angle, reflecting a tuning that was better adapted for the perturbed visuomotor environment.

Having established a within-trial interpretation for each component, we next examined across-trial learning dynamics. For visualization purposes, we gently smoothed all TCA trial factors by a Gaussian filter with a standard deviation of 1.5 trials. Across all reach angles, the *early component* was typically flat and insensitive to the visuomotor perturbation (fig. 7d, left). In contrast, the *learned component* activated soon after the perturbation was applied, although the rapidness of this onset varied across reach angles (fig. 7d, right). Together, this reinforces our earlier observation that adaptation to the visuomotor rotation typically involves the production of new neural dynamics (captured by the *learned component*), rather than the suppression of maladaptive dynamics (captured by the *early component*).

Finally, the *corrective component* was consistently correlated with the animal’s behavioral performance on all reach angles (*p <* 0.05, Spearman’s rho test). Since performance differed across reach angles, we separately plotted the *corrective component* (blue) against the time to acquire the target (black) for each reach angle (fig. 7e). Remarkably, in many cases, the corrective component provided an accurate trial-by-trial prediction of the reach duration, meaning that trials with a large corrective movement took longer to execute.

Together, these results demonstrate that TCA can identify, in a purely unsupervised manner, both learning dynamics across trials and single trial neural dynamics. Indeed, each trial factor can be related to within-trial behaviors, such as error-prone cursor movements and their subsequent correction. Furthermore, these basic interpretations largely replicate across all eight reach angles, despite differences in the learning rate within each of these conditions. Most intriguingly, a *single* trial factor, extracted only from neural data, can directly predict execution time on a trial by trial basis, without ever having direct access to this aspect of behavior (fig. 7e).

## 3 Discussion

Recent experimental technologies enable us to record from more neurons, at higher temporal precision, and for much longer time periods than ever before [4, 8, 11], thereby simultaneously increasing the size and complexity of datasets along three distinct modes. However, methods for multi-timescale dimensionality reduction that describe both rapid neural dynamics within trials and long-term changes in neural dynamics across-trials are still lacking. As a result, experimental investigations of neural circuits are often confined to a single timescale, even though bridging our understanding across multiple timescales is of great interest [9]. Here we demonstrated a unified approach, TCA, that simultaneously recovers low-dimensional and interpretable structure across neurons, time within trials, and trials.

TCA and other tensor decomposition techniques have been extensively studied from a theoretical perspective [29, 63–65], and have been applied to a variety of biomedical problems [66–68]. Several studies have applied tensor decompositions to EEG and fMRI data, most typically to model differences across subjects or Fourier/wavelet transformed signals [69–72], rather than across trials [73]. A recent study examined trial-averaged neural data across multiple neurons, conditions, and time within trials as a tensor, but they did not study trial-to-trial variability, and only examined different unfoldings of the data tensor into matrices, rather than applying TCA directly to the data tensor [74]. Other studies have modeled the receptive fields of neurons in auditory and visual cortex as third-order tensors with low-rank structure [75, 76]. We go beyond these previous studies by applying TCA to a broader class of artificial and experimental datasets, drawing a novel connection between TCA and theories of gain modulation, and demonstrating that visualization and analysis of the TCA trial factors can directly yield functional clustering of neural populations (i.e., cell assemblies) as well reveal learning dynamics on trial-by-trial basis.

In particular, we demonstrated that TCA reveals a simple description of learning in an artificial nonlinear neural network trained to solve the analog of a motion discrimination task [32]. TCA discovered a one-dimensional learning process in which initial, small random selectivity is monotonically amplified over time to yield the final learned decision making dynamics. Moreover, cell-type information extracted by TCA in an unsupervised manner enabled us to re-organize the network’s connectome, thereby yielding conceptual insights into how this connectome gives rise to mechanisms for decision making. Also, in calcium imaging data recorded from rodent pre-frontal cortex during a maze navigation task, TCA uncovered functional subsets of neurons that fired sequentially within trials, and whose amplitude on each trial selectively mapped onto task-relevant variables, including starting location, ending location and reward (in that order).

Finally, in electrophysiological recordings from macaque motor and premotor cortex, TCA revealed a simple two-dimensional learning process in response to a BMI perturbation. Interestingly, this learning process did not involve extinguishing maladaptive dynamics that were established in the pre-perturbation period (i.e., the “early component” identified by TCA). Rather, it involved the addition of two components that compensated for the maladaptive dynamics. The first “corrective” component was a suboptimal, late stage within-trial correction that was active in trials soon after the perturbation, which extinguished over trials to give rise to a second “learned” component that implemented a more optimal early stage within-trial correction. Moreover, the late stage correction could predict time to target acquisition on a trial-by-trial basis. Importantly, all of these results were discovered purely from the neural data and not behavioral measurements, suggesting that TCA can uncover unexpected and otherwise unobservable neural dynamics in a data-driven, unsupervised manner.

In addition to the empirical success of TCA in diverse scenarios presented here, there are three other reasons we expect TCA to have widespread utility in neuroscience. First, TCA is arguably the simplest generalization of PCA that can handle trial-to-trial variability. Given the widespread adoption of PCA, we believe that TCA may also enjoy widespread adoption and success, especially as technologies enabling long-term and large-scale recordings become more accessible. Second, as we have shown, TCA has an intriguing interpretation as a network model with low-dimensional gain-modulated inputs. This model is supported by experimental evidence in many contexts [22, 37, 76–78], and underlies influential theories of cortical computation [30, 31] and perceptual learning [79].

Third, while TCA is a simple generalization of PCA, its theoretical properties are strikingly more favorable. A fundamental limitation of PCA is that the components it recovers are restricted to be orthogonal to each other, and moreover these components can be rotated amongst each other without changing the reconstruction error. This invariance to rotations in PCA leads to a fundamental ambiguity, and so the factors identified by PCA are unlikely to be directly interpretable as biological signals (see *Methods*, section 4.4.2). In contrast, the factors identified by

TCA are not invariant to many transformations [29], yielding more interpretable results. This advantage was first demonstrated in fig. 2 where the factors recovered from neural firing rates, matched the underlying parameters of the model neural network in a one-to-one fashion. Similarly, in the rodent prefrontal analysis, TCA uncovers demixed factors that individually correlate with interpretable task variables, whereas PCA does not (compare fig. 5 to fig. 5 supp. 1). And finally, when applied to neural activity during BMI learning, TCA consistently found, across multiple reach angles, a “corrective factor” that significantly correlated with behavioral performance on a trial-by-trial basis (fig. 7).

In this paper, we examined the simplest form of TCA by making no assumptions about the temporal dynamics of neural activity within trials or the dynamics of learning across trials. As a result, we obtain extreme flexibility: for example, trial factors could be discretely activated or inactivated on each trial (fig. 5), or they might emerge incrementally over longer timescales (fig. 6). However, future work could augment TCA with additional structure and assumptions, such as a smoothness penalty or dynamical systems structure within trials [16]. Intriguingly, a dynamical system could just as easily be incorporated along the trials axis of the data tensor to potentially relate high-dimensional neural activity to low-dimensional models of learning [80].

Further work in this direction could connect TCA to a large body of work on fitting latent dynamical systems to reproduce within-trial firing patterns. In particular, single trial neural activity has been modeled with linear dynamics [81–84], switched linear dynamics [85, 86], linear dynamics with nonlinear observations [17], and nonlinear dynamics [18, 87]. In practice, these methods require many modeling choices, validation procedures, and post-hoc analyses. Simple linear models have a relatively constrained dynamical repertoire [12], while models with nonlinear elements often have greater predictive abilities [17, 18], but at the expense of interpretability. In all cases, the learned representation of each trial (e.g., the initial condition to a nonlinear dynamical system) is not transparently related to single trial data. In contrast, the trial factors identified by TCA have an extremely simple interpretation as introducing trial-specific linear gain modulation. Overall, we view TCA as a simple and complementary technique to identifying a full dynamical model, as has been previously suggested for PCA [12].

An important property of TCA is that it extracts salient features of a dataset in a data-driven, unbiased fashion. Such unsupervised methods are a critical counterpart to supervised methods, such as regression, which can directly assess whether a dependent variable of interest is represented in population activity. Recently developed methods like *demixed PCA* [34] combine regression with dimensionality reduction to isolate linear subspaces that selectively code for variables of interest. Again, we view TCA as a complementary approach, with at least three points of difference. First, like trial-concatenated PCA and GPFA, demixed PCA only reduces dimensionality within trials by identifying a different low-dimensional temporal trajectory for each trial. In contrast, TCA identifies a common low-dimensional temporal trajectory (temporal factors) for all trials, which are modulated by different amplitudes (trial factors) on each trial. Second, demixed PCA can separate neural dynamics in cases where trials have discrete conditions and labels, such as in the rodent prefrontal analysis in fig. 5; however, it is not designed to handle continuous dependent variables, such as those describing learning dynamics (see fig. 3 and fig. 7). Furthermore, unsupervised techniques like TCA can identify unexpected cognitive states and dynamics corresponding to unknown or difficult to measure dependent variables. Finally, the same rotation invariance of PCA is present within the linear subspaces identified by demixed PCA. Thus, both PCA and demixed PCA are fundamentally subspace identification algorithms, while TCA can often extract directly meaningful features from data, such as clusters of functional cell types or neural populations that grow or shrink in magnitude across trials.

An intriguing direction for future research is to expand TCA to higher-order tensors beyond those encoding neurons, timepoints, and trials. For example, we can also record across multiple subjects learning to solve the same task, yielding a fourth order data tensor, with individual subjects as the fourth index. Similarly, if an individual subject is taught multiple learning tasks, one could encode experimental task or condition as a fourth index. However, directly applying TCA to these tensors may be undesirable, since we record from different neural populations in different subjects and the learning rate may vary from subject-to-subject or from task-to-task. Instead, we could model such data via coupled tensor factorizations [88] which allow some measured tensor axes to be fit as common factors, while others are fit in a separate and unconstrained fashion. For instance, we could assign separate neuron and trial factors for each subject, but use shared temporal factors across subjects if they are hypothesized to share similar low-dimensional within trial cognitive dynamics. This scheme could extract common circuit dynamics from small numbers of neurons through increased statistical power obtained via pooling across multiple subjects. Moreover, the separate neuron factors would then provide “translations” between subjects, by revealing how the same cognitive variable is encoded in different population activity patterns in different subjects. In essence, while moving from second to third-order tensor methods provides a new window into how circuit dynamics changes across trials to mediate learning, moving additionally to fourth order tensor methods may provide new insights into how the learning dynamics itself changes across subjects and tasks.

Overall, this work highlights the prevalence of tensor structure in neural datasets and demonstrates that exploiting this structure can provide extremely useful insights into complex, multi-timescale, high-dimensional neural data, including the unsupervised discovery of cell assemblies, within trial neural dynamics underlying perceptions, actions and thoughts, and across trial learning dynamics. Just as PCA has become part of the standard canon of neural data analyses for trial-averaged neural recordings, the combined simplicity and power of TCA suggests it may have widespread utility in the analysis of multineuronal data at the level of single trials.

## 4 Methods

### 4.1 Key Resources Table

### 4.2 Contact for Reagent and Resource Sharing

Further requests for resources should be directed to and will be fulfilled by the Lead Contact, Alex H. Williams

### 4.3 Data and Software Availability

We provide specialized tools for fitting and visualizing TCA in https://github.com/ahwillia/tensortools. Other resources for fitting tensor decompositions include [93–95].

| Reagent or Resource | Source | Identifier |
| --- | --- | --- |
| Software and Algorithms |  |  |
| tensortools | This paper | https://github.com/ahwillia/tensortools |
| Alternating Least Squares | [27] | N/A |
| SciPy | [89] | https://scipy.org/ |
| Matplotlib | [90] | https://matplotlib.org/ |
| scikit-learn | [91] | http://scikit-learn.org/ |
| PyTorch | none | http://pytorch.org/ |
| MATLAB | MathWorks | https://www.mathworks.com/products/matlab.html |
| Simulink Realtime | MathWorks | https://www.mathworks.com/products/simulink-real-time.html |
| Experimental Models: Organisms/Strains |  |  |
| C57BL/6J mice | The Jackson Laboratory | 000664 |
| Rhesus macaque (*Mucacca Mulatta*) | Wisconsin and Yerkes primate centers | N/A |
| Recombinant DNA |  |  |
| pGP-CMV-GCAMP6m | [92] | #40754, https://www.addgene.org/Douglas_Kim/ |
| Other |  |  |
| Miniature fluorescence microscope | Inscopix | https://www.inscopix.com/nvista |
| Utah Microelectrode Arrays | Blackrock Microsystems | http://blackrockmicro.com/neuroscience-research-products/low-noise-ephys-electrodes/blackrock-utah-array/ |
| Cerebus System | Blackrock Microsystems | http://blackrockmicro.com/neuroscience-research-products/neural-data-acquisition-systems/cerebus-daq-system/ |

### 4.4 Method Details

#### 4.4.1 Notation and Terminology

Colloquially, a tensor is a data array or table with multiple axes or dimensions. More formally, the axes are called *modes* of the tensor, while the *dimensions* of the tensor are the lengths of each mode. Throughout this paper we consider a tensor with three modes with dimensions *N* (number of neurons), *T* (number of timepoints in a trial), and *K* (number of trials).

The number of modes is called the *order* of the tensor. We denote vectors (order-one tensors) with lowercase boldface letters, e.g., **x**. We denote matrices (order-two tensors) with uppercase boldface letters, e.g., **X**. We denote higher-order tensors (order-three and higher) with boldface calligraphic letters, e.g., 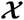. Scalars are denoted by non-boldface letters, e.g., *x* or *X*. We use **X**^*T*^ to denote the transpose of **X**. We aim to keep other notation light and introduce as it is first used — readers may refer to [33] for notational conventions.

#### 4.4.2 Matrix and Tensor models

Neural population activity is commonly represented as a matrix with each row holding a neuron’s activity trace [12]. Let **X** denote an *N × T* matrix dataset in which *N* neurons are recorded over *T* time steps. For spiking data, **X** may denote trial-averaged spike counts or a single-trial spike train smoothed with a Gaussian filter. If fluorescence microscopy is used in conjunction with voltage or calcium indicators, the data entries could be normalized fluorescence (∆F*/*F).

PCA is a special case of *matrix decomposition*. A matrix decomposition model approximates the data **X** as a rank-*R* matrix, 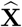, yielding *R* components. This approximation can be expressed as the product of an *N × R* matrix **W** and a *T × R* matrix **B**:

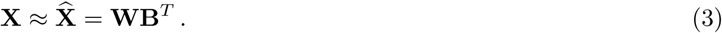

We call the columns of **W** neuron factors, denoted **w**^*r*^, and the columns of **B** temporal factors, denoted **b**^*r*^. The rows of **W**, denoted **w**_*n*_, provide an *R*-dimensional description of each neuron’s activity trace. Likewise the rows of **B**, denoted **b**_*t*_, provide an *R*-dimensional description of the full neural population activity pattern at each timepoint. In order to reduce the dimensionality of the data we chose *R < N* and *R < T*. Note that eq. (3) is equivalent to eq. (1) in the *Results*.

Perhaps the simplest matrix decomposition problem is to identify a rank-*R* decomposition that minimizes the squared reconstruction error:

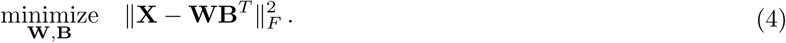

Here, 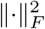 denotes the squared *Frobenius norm* of a matrix, which is simply the sum of squared matrix elements:

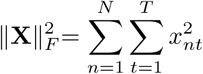

PCA provides *one* solution to eq. (4). Most critically, the PCA solution constrains the neuron factors and temporal factors to be orthogonal, meaning that **W**^*T*^ **W** and **B**^*T*^ **B** are diagonal matrices. However, this solution does not uniquely minimize the squared reconstruction error. In fact, there is a continuous manifold of matrix decompositions that solve eq. (4), since any invertible linear transformation **F** can produce a new set of parameters, **W**′ = **WF**^−1^ and **B**′ = **BF**^*T*^ that produce an equivalent reconstruction of the data:

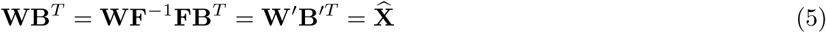

This result — sometimes called the *rotation problem* — has a fundamental consequence: if the data were truly generated as a combination of *R* low-dimensional components, then PCA *cannot not recover these ground truth components*. At best, PCA can only be expected to recover the same linear subspace of the true components.

In essence, after fitting a PCA model, one might be tempted to interpret the columns of **W** as identifying subpopulations of neurons with firing patterns given by the columns in **B**. However, eq. (5) shows that these putative sub-populations can be linearly mixed by a broad class of transformations, so long as the components are mixed by the appropriate inverse transformation. Thus, the latent factors identified by PCA are poorly constrained, and it is better to interpret PCA as finding an orthogonal coordinate basis for visualizing data. As reviewed below, the optimization problem addressed by TCA has superior uniqueness properties relative to eq. (4), which gives us greater license to directly interpret the TCA factors as potentially biologically meaningful neural populations and activity patterns.

TCA is a natural generalization of PCA to higher-order tensors. Let 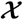 denote a *N* × *T* × *K* data tensor, and let *x*_*ntk*_ represent the activity of neuron *n* at time *t* on trial *k*. For a third-order tensor, TCA finds a set of three factor matrices, **W**, **B**, and **A**, with dimensions *N × R*, *T × R*, and *K × R*, respectively. As before, the columns of **W** are the neuron factors, the columns of **B** are the temporal factors. Analogously, the columns of **A** are the trial factors, denoted **a**^*r*^, and the rows of **A**, denoted **a**_*k*_, embed each trial into an *R*-dimensional space.

To reformulate eq. (2) into an equivalent matrix equation, let **X**_*k*_ denote an *N × T* matrix holding the data from trial *k*. TCA models each trial of neural data as:

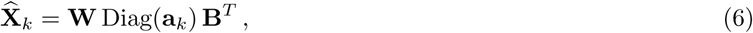

where Diag(**a**_*k*_) embeds **a**_*k*_ as the diagonal entries of an *R × R* matrix. Again, eq. (6) is equivalent to eq. (2) in the *Results*. In this paper, we also employed the *nonnegative* TCA model, which simply adds a constraint that all factor matrices have nonnegative elements:

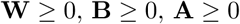

Nonnegative TCA has been previously studied in the tensor decomposition literature [64, 96–98], and is a higher-order generalization of nonnegative matrix factorization (NNMF) [60, 99]. Similar to eq. (3), in this paper both unconstrained and nonnegative TCA were fit to minimize the squared reconstruction error:

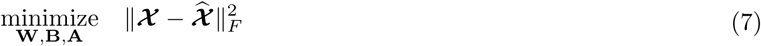

Both PCA and TCA can be extended to incorporate different loss functions, such as a Poisson negative log-likelihood [100], however we do not consider these models in this paper.

Fitting TCA to data is a nonconvex problem. Unlike PCA, there is no efficient procedure for achieving a certifiably optimal solution [65]. We use established optimization algorithms to minimize eq. (7) from an initial guess (see section 4.4.3). Although this approach may converge to local minima in the objective function, our results empirically suggest that this is not a major practical concern. Indeed, as long we does not choose too many factors (too large an *R*) and use nonnegative factors, we find that the multiple local minima yield similar parameter values and similar reconstruction error.

An important advantage of TCA is that the low-dimensional components it uncovers are often “essentially unique,” up to permutations and scalings. More precisely, in [29], it was proven that every local minimum of the TCA objective function is isolated in parameter space; it is not part of a continuous manifold of parameters that achieve *exactly* the same reconstruction error, as in matrix factorization described above. Instead, this continuous degeneracy, or ambiguity is replaced by a much more benign ambiguity, namely a set of solutions with the same reconstruction error related to each other simply by permutations and rescalings. For instance, the columns of **W**, **B**, and **A** can be jointly permuted without affecting the model. Also, the columns of any pair of **W**, **B**, and **A** can be jointly rescaled. For example, if the *r*^th^ column of **W** is multiplied by a scalar *s*, then the *r*^th^ column of either **B** or **A** can be divided by *s* without affecting the model’s prediction. These transformations, which are also present in PCA, are inconsequential since the direction of the latent factors and total size of any set of factors, rather than their order, are of primary interest. Thus the parameter set corresponding to the global minimum of TCA is essentially unique, up to permutations and scalings. Of course, in general we are not guaranteed to find this global minimum, but as we have shown in the main text, in situations where we do not choose too many factors, all the local minima we find using multiple runs of TCA achieve similarly low reconstruction error, and moreover are close to each other in parameter space. In such a situation, all the local minima likely cluster near the global minimum, and the resultant parameter values are likely to be biologically meaningful, or interpretable.

In summary, when the factors are all linearly independent (i.e., **W**, **B**, and **A** have full column rank), TCA is, in the sense described above, provably unique up to rescalings and permutations [29]. TCA can nevertheless be difficult to optimize if latent factors are approximately linearly dependent [101]. To quantify and monitor this possibility, we computed a similarity score between TCA models based on the angles between the extracted factors (see section 4.5.1). In practice, we did not find this to be a critical problem.

#### 4.4.3 Model optimization

TCA can be applied to neural data by a series of simple steps (Figure 2f). First, to incorporate the common assumption that latent neural firing rates are smooth in time [16], spiking data can be temporally smoothed (e.g., with a Gaussian filter). The width of this smoothing filter affects the smoothness of the latent temporal factors recovered by TCA. Analogous smoothness hyperparameters are present in other dimensionality reduction methods. For example, in GPFA, the timescale of latent dynamics are set by the autocorrelation in the prior’s covariance matrix [16]. Depending on the dataset, it may be important to apply other common preprocessing steps, such as z-scoring the activity traces of neurons, or applying variance-stabilizing transformations such as taking the square root of spike counts [12].

Like many dimensionality reduction methods, TCA can only be fit by iterative optimization algorithms. While these procedures may get stuck in sub-optimal local minima, in practice we found that all optimization fits converged to similar reconstruction errors. Other techniques, such as nonnegative matrix factorization [60], also demonstrate practical success while being NP-hard in terms of worst-case analysis [102].

Specialized algorithms for fitting TCA are an area of active research. We used the classic method of *alternating least-squares* (ALS) to obtain estimates of the factor matrices. ALS is motivated by the observation that fixing two of the factor matrices and optimizing over the third in eq. (7) is a least-squares subproblem that is convex and has a closed-form solution. For illustration, consider optimizing the neuron factors **W**, while temporarily fixing the within-trial factors, **B**, and the trial factors **A**. This yields the following update rule:

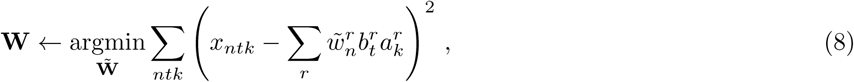

which can be solved as a linear least-squares matrix problem. In particular, with some manipulation of the indices, eq. (8) can be rearranged into a matrix equation (see [33]) and solved by standard matrix library routines. This procedure is then cyclically repeated: the temporal factors **B** are updated while fixing **W** and **A**, then the trial factors **A** are updated while fixing **W** and **B** and so on until the objective function converges. The ALS algorithm is available in several open-source packages [**tensortoolbox2.6**, 94, 95], and is reviewed in [33]. For nonnegative TCA, we solved each sub-problem using a specialized nonnegative least squares solver [103], instead of standard least-squares.

#### 4.4.4 Linear gain-modulated model network

In fig. 2, we constructed a linear network model with three input neurons connected to *N* = 50 observed neurons by random Gaussian weights. The outgoing weights of each input neuron were normalized to unit Euclidean length. Each input neuron had a different temporal firing pattern lasting *T* = 150 time steps, parameterized as probability density functions of Gamma distributions. The trial-specific amplitude of the first two input neurons were respectively parameterized as increasing and decreasing logarithmically spaced points over *K* = 100 trials. The amplitude of the third input neuron linearly increased for *K <* 50 and then linearly decreased to the same starting value. All within-trial waveforms and across-trial amplitude vectors were normalized to unit Euclidean length. As described in the *Results*, the activity of all neurons is modeled by the same equations as TCA (eq. (2)). Independent and identically distributed Gaussian noise with a standard deviation of 0.01 was added to the simulated data. ICA and PCA were performed on this simulated dataset via the scikit-learn Python package [91].

#### 4.4.5 Nonlinear recurrent neural network model

We simulated a discrete-time recurrent neural network with a hyperbolic tangent nonlinearity.

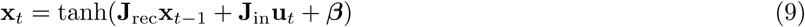

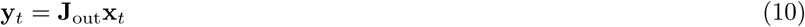

Here, **x**_*t*_ is a vector of *N* neural firing rates of the recurrently connected neural population at time *t*, **u**_*t*_ and **y**_*t*_ are the inputs and outputs of the network, **J**_rec_, **J**_in_, **J**_out_ are synaptic weight matrices for the recurrent, input, and output connections, and ***β*** is a *N*-dimensional vector of bias terms. The input and output of the were one-dimensional signals, as illustrated in fig. 3a. Thus, the recurrent synaptic weights were held in a *N × N* matrix, **J**_rec_, the input weights were held in a *N ×* 1 matrix, **J**_in_, and the output weights were held in a *N ×* 1 matrix, **J**_out_.

On each trial, the input signal to the network consisted of *T* = 40 independent draws from a standard normal distribution with mean *µ* = 1 or *µ* = *−*1 (chosen randomly with equal probability on each trial). The goal of the network was to produce a positive output (*y_t_ >* 0) when the input was net-positive, and produce a negative output (*y_t_ <* 0) when the input was net-negative. The performance of the network on each trial was measured by a logistic loss function (applied to the output on the final time step, *y*_*T*_):

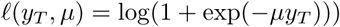

For each simulated trial, we used the deep learning framework PyTorch to compute the gradient of this loss function with respect to all network parameters {**J**_rec_, **J**_in_, **J**_out_, **β**} via the backpropagation through time algorithm. A small parameter update in the direction of the negative gradient for each weight matrix was applied after each trial (stochastic gradient descent, with a learning rate of 0.005). This was repeated for *K* = 750 trials. The activity of the recurrent units (**x**_*t*_ in eq. (9)) over all timepoints and trials was collected into a *N × T × K* tensor for analysis.

#### 4.4.6 Mouse spatial navigation task

We injected 500 nL of AAV2/5-CaMKII*α*-GCaMP6m into the medial prefrontal cortex (AP: 1.9, ML: 0.95, DV: 2.25, relative to bregma) into mice aged ∼8 weeks. Approximately one week after virus injection, we installed glass-bottom stainless steel guide tubes into the prefrontal cortex to enable deep brain optical imaging using a 1 mm diameter GRIN microendoscope (1050-002176, Inscopix). Two weeks following guide tube surgery, we checked for cellular Ca^2+^ signals with a miniaturized fluorescence microscope (nVista HD, Inscopix). Animals with robust Ca^2+^ responses were selected for further behavioral study. Mice selected for behavioral training underwent water restriction (1 mL per day) to reach ∼85% of their *ad libitum* weight.

Mice performed spatial navigation on a custom-built elevated plus maze. The center-to-end arm length of the maze was 38 cm. By blocking one of the arms with an opaque barrier, the plus maze could be converted into a T-maze with any of the four arms as the stem. Additional gates on each of the arms (at ∼ 15 cm from the end) could be used to confine the mouse at the arms. At the end of each arm, a proximity sensor enabled detection of the mouse and a water spout allowed for reward delivery. The maze was placed in a rectangular housing whose four side walls were uniquely defined by distinctly patterned curtains.

The mice performed 100-150 trials on each session. At the beginning of each trial, the experimenter placed the mouse in the stem arm of the T-maze with the corresponding gate closed. After 5 s holding time in the stem arm, the “start” gate was opened to allow the mouse to run to either end of the T-maze. Once the mouse was detected in one of the ends, the “end” gate was closed behind the mouse to confine it in the chosen arm for another 5 s. If the mouse’s choice was consistent with the reward contingency, 5-10 *µ*L of water was delivered to the spout. Trained mice typically made the run in 2 s; hence the typical trial was ∼12 s long. At the end of each trial, the experimenter retrieved the mouse and wiped the maze with ethanol.

During trials, we recorded prefrontal Ca^2+^ activity at 20 Hz using the miniature fluorescence microscope. An overhead camera (DMK 23FV024, The Imaging Source) mounted above the behavioral apparatus synchronously recorded the position of the mouse on the maze. To extract cells and their activity traces from the Ca^2+^ movies, we followed a procedure previously described in [5], and we then tracked individual neurons across sessions using previously described methods [104].

The tensor representation of neural activity requires that the number of samples within each trial be the same for all trials, whereas the mice took a variable amount of time to complete each trial. Hence, we used the largest number of intra-trial samples common to all trials (or, equivalently, the duration of the shortest trial) as the length of the intra-trial time dimension. We chose to temporally align trials to the end of each trial, because the mice showed more consistent behavior across trials at the ends (i.e. approaching the choice arm and consuming reward, if available) rather than the beginnings (where mice could take variable time to initiate motion after opening of the start gate).

Along the trial dimension of the tensor, we simply concatenated trials across days. However, all Ca^2+^ activity traces were normalized to the range [0, 1] based on the cell’s minimum and maximum fluorescence values on each day. This normalization procedure was crucial for forming across-day tensors, since the exact amplitude of a Ca^2+^ trace was dependent on precise, micron-level axial positioning of the microscope — which could vary randomly from session to session.

#### 4.4.7 Primate BMI task

The monkey’s hands were restrained for the full duration of the experiment. Voltage signals were band-pass filtered from each electrode (250 Hz - 7.5 KHz). A spike was recorded whenever these filtered signals crossed below a threshold of −4.5 times the root-mean-square voltage.

The neural recordings from PMd and M1 were used jointly and without distinction to train a BMI decoder by the recalibrated feedback-intention trained Kalman filter (ReFIT) procedure [61]. At the start of each session, the monkey observed 600 trials of automated cursor movements from the center of the workspace to one of 8 radially arranged targets at a distance of 12 cm. During these observation trials, the cursor velocity began at 8 cm/s, and increased by 2 cm/s every 200 trials. Under the premise that the monkey is imagining the intended task during these observation trials, we used the neural activity and cursor kinematics to fit a Kalman filter decoder. The velocity gain of the decoder was calibrated by the experimenter to help the monkey achieve fast reaches (improved by high gain) while still holding the cursor steady (improved by low gain).

The monkey then executed instructed-delay cursor movements to indicated radial target locations, before returning to the center position and repeating the cursor movement to another target. This essential behavioral paradigm has been previously described [105]. Each target position and the center position were indicated on the screen. Monkeys started by holding the cursor on the central target continuously for 500 ms. After a randomized delay (sampled uniformly from 400-800 ms), monkeys moved the cursor within a 4 × 4 cm acceptance window of the cued target. This target also had to be held continuously for 500 ms. The target changed color to signify the hold period. If the cursor left the acceptance window, the timer was reset, but the trial was not immediately failed. Monkeys had 2 s to acquire the target. Success was accompanied with a liquid reward, along with a success tone. Failure resulted in no reward, and a failure tone. The center target was then presented, which the monkeys also had to acquire and hold.

For our analysis, we collected the non-sorted spiking activity of all *N* = 192 multiunit recordings during all center to outward cursor reaches (reaches back to the center were not analyzed). Spike times were aligned to the end of the delay period (*t* = 0) and ended at the time of first target acquisition or after two seconds had elapsed and the target was still not required. The data tensor was zero padded to ensure a consistent trial length of two seconds. Data were smoothed within each trial with a Gaussian filter with a standard deviation of 50 ms (same as in [34]). Using a smaller filter did not qualitatively effect the trial factors extracted by TCA, but resulted in less smooth temporal factors.

### 4.5 Quantification and Statistical Analysis

#### 4.5.1 TCA model analysis

Unlike PCA (but similar to ICA and other methods), TCA needs to be iteratively optimized to minimize a cost function. In theory, each optimization run may converge to a sub-optimal local minimum. Additionally, the number of components in the model can affect the final result [63]. This is different from PCA where the largest components do not change by adding additional components (a consequence of the Eckert-Young theorem; [106]). Thus, we fit all TCA models from multiple initial parameters and with different numbers of low-dimensional factors. We then inspect this ensemble of models for a consistent and interpretable summary of the data.

The most basic metric to compare models is the squared reconstruction error, since this is what TCA aims to minimize. For interpretability, we normalize the reconstruction error on a scale of zero to one:

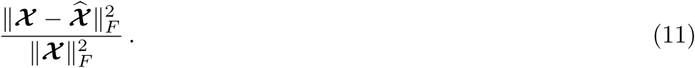

We typically visualize reconstruction error as a function of the number of model components (see, e.g., fig. 4g), which we call an “error plot.”

As discussed in section 4.4.2, TCA is invariant to permutations and rescalings of the factors. In PCA, the components are often normalized to unit Euclidean length and ordered by variance explained. An analogous procedure exists for TCA [33]. First, rescale the columns of **W**, **B**, and **A** to be unit length, and absorb these scalings into *λ*_*r*_ for each component *r*. Then the estimate of the data becomes:

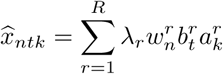

If desired, the components can be sorted by decreasing *λ*_*r*_.

To quantify the similarity of two fitted TCA models, we used a similarity score based on the angles between latent factors [107]. Formally, for two TCA models, {**W**, **B**, **A**} and {**W**′, **B**′, **A**′}, the similarity score is:

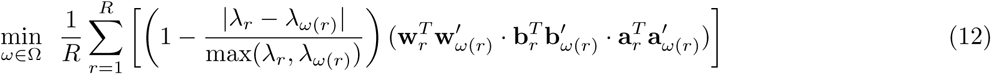

Where Ω denotes the set of all permutations of the factors, and *ω* is a particular permuation. For example, for a three component model (*R* = 3) the score is computed for all possible permutations, *ω* = {1, 2, 3}, {2, 1, 3}, {3, 2, 1}, and {3, 1, 2}, and the lowest score is taken. For TCA models with more than 10 components, enumerating all permutations can be computationally prohibitive. In these cases we match factors in a greedy fashion to identify a permutation that provides a good (though not certifiably optimal) alignment of the models. Note that this measurement of model similarity is quite severe, since the distance of each pair of factors are multiplied – if any single dimension is orthogonal, For our datasets, models with similarity scores above 0.8 were qualitatively similar and led to similar quantitative results in post-hoc analyses. Models with similarity scores within the 0.6−0.8 range also appeared quite similar in our applications.

#### 4.5.2 Mouse spatial navigation task

We quantified the *dimensionality* of a single neuron across trials by the following quantity:

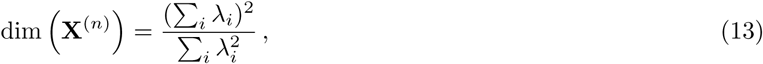

where *λ*_*i*_ are the eigenvalues of the covariance matrix; i.e., 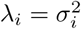 where *σ*_*i*_ are the singular values of **X**^(*n*)^, which is a *K × T* matrix holding the activity of neuron *n* across all trials. This is a continuous measure of dimensionality used in condensed matter physics, and was previously applied to analyze neural circuits [108]. For example, (13) reduces to *N* when all the *λ*_*i*_ are evenly distributed and take the same value, and reduces to 1 if only one *λ*_*i*_ is nonzero. For uneven distributions of *λ*_*i*_, this measure sensibly interpolates between these two extremes.

#### 4.5.3 Primate BMI task

In fig. 7, statistical tests on the mean preferred angle of TCA components were performed using PyCircStat (https://github.com/circstat/pycircstat). Statistical tests on Spearman’s rho were computed using the SciPy statistics module (https://docs.scipy.org/doc/scipy-0.14.0/reference/stats.html).

### 4.6 Experimental model and subject details

#### 4.6.1 Mice

The Stanford Administrative Panel on Laboratory Animal Care approved all mouse procedures. We used male C57BL/6 mice, aged ∼8 weeks at start. Throughout the entire protocol, we monitored the weight daily and looked for signs of distress (e.g., unkempt fur, hunched posture). Mice were habituated to experimenter handling and the behavioral apparatus for ∼2 weeks prior to the five day behavioral protocol.

#### 4.6.2 Monkey

Recordings were made from motor cortical areas of an adult male monkey, R (*Macaca mulatta*, 15 kg, 12 years old), performing an instructed delay cursor task. The monkey had two chronic 96-electrode arrays (1 mm electrodes, spaced 400 *µ*m apart, Blackrock Microsystems), one implanted in the dorsal aspect of the premotor cortex (PMd) and one implanted in the primary motor cortex (M1). The arrays were implanted 5 years prior to these experiments. Animal protocols were approved by the Stanford University Institutional Animal Care and Use Committee.

## Acknowledgments

The authors thank Jeff Seely (Cognescent Corporation) and Casey Battaglino (Georgia Tech) for discussions pertaining to this work. A.H.W. was supported by the Department of Energy Computational Science Graduate Fellowship program. T.H.K. was supported by a Stanford Graduate Fellowship in Science & Engineering. F.W. was supported by a National Science Foundation Graduate Research Fellowship. S.V. was supported by a National Science Foundation Graduate Research Fellowship, a Ric Weiland Stanford Graduate Fellowship, National Institutes of Health F31 training grant, and the Stanford Center for Mind, Brain and Computation. K.V.S. was supported by the US National Institutes of Health Director’s Pioneer Award 8DP1HD075623, US National Institutes of Health Director’s Transformative Research Award (TR01) from the NIMH #5R01MH09964703, Defense Advanced Research Projects Agency NeuroFAST award from BTO #W911NF-14-2-0013, the Simons Foundation, and the Howard Hughes Medical Institute. M.S. was supported by the National Institutes of Health (#1R21NS104833-01), the National Science Foundation (#1707261), and the Howard Hughes Medical Institute. Work by T.G.K. was supported by the U.S. Department of Energy, Office of Science, Office of Advanced Scientific Computing Research, Applied Mathematics program in a grant to Sandia National Laboratories, a multimission laboratory managed and operated by National Technology and Engineering Solutions of Sandia, LLC., a wholly owned subsidiary of Honeywell International, Inc., for the U.S. Department of Energy’s National Nuclear Security Administration under contract DE-NA-0003525. S.G. was supported by the Burroughs Wellcome Foundation, the McKnight Foundation, the James S. McDonnnell Foundation, the Simons Foundation, and the Office of Naval Research.

## Figure Supplements

**Figure 2, Supplement 1.**
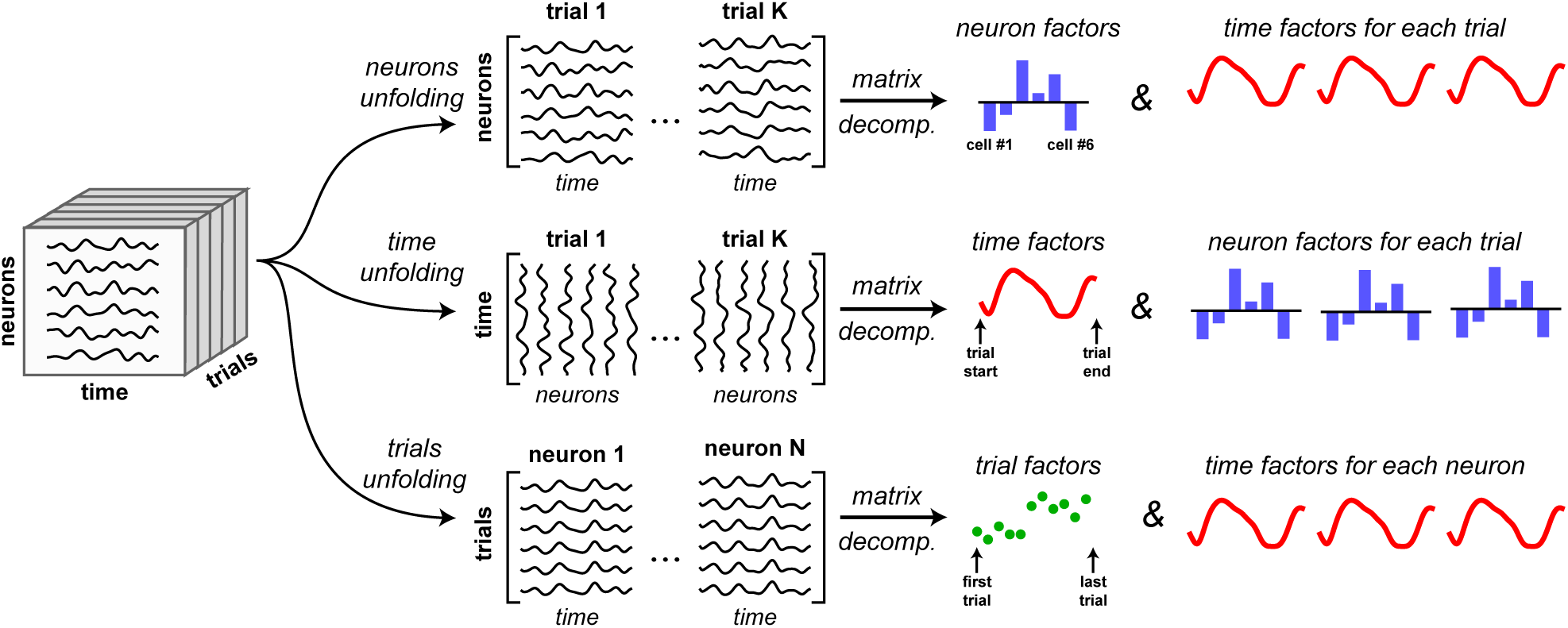
Illustration of *tensor unfolding* for applying matrix decompositions to tensor datasets. A *N × T × K* dimensional tensor can be reshaped into three different matrices: a “neurons unfolding” with dimensions *N × T K*, a “time unfolding” with dimensions *T × NK*, and a “trials unfolding” with dimensions *K × NT*. Applying PCA or other matrix decomposition methods to each unfolding yields a different set of low-dimensional factors.

**Figure 3, Supplement 1.**
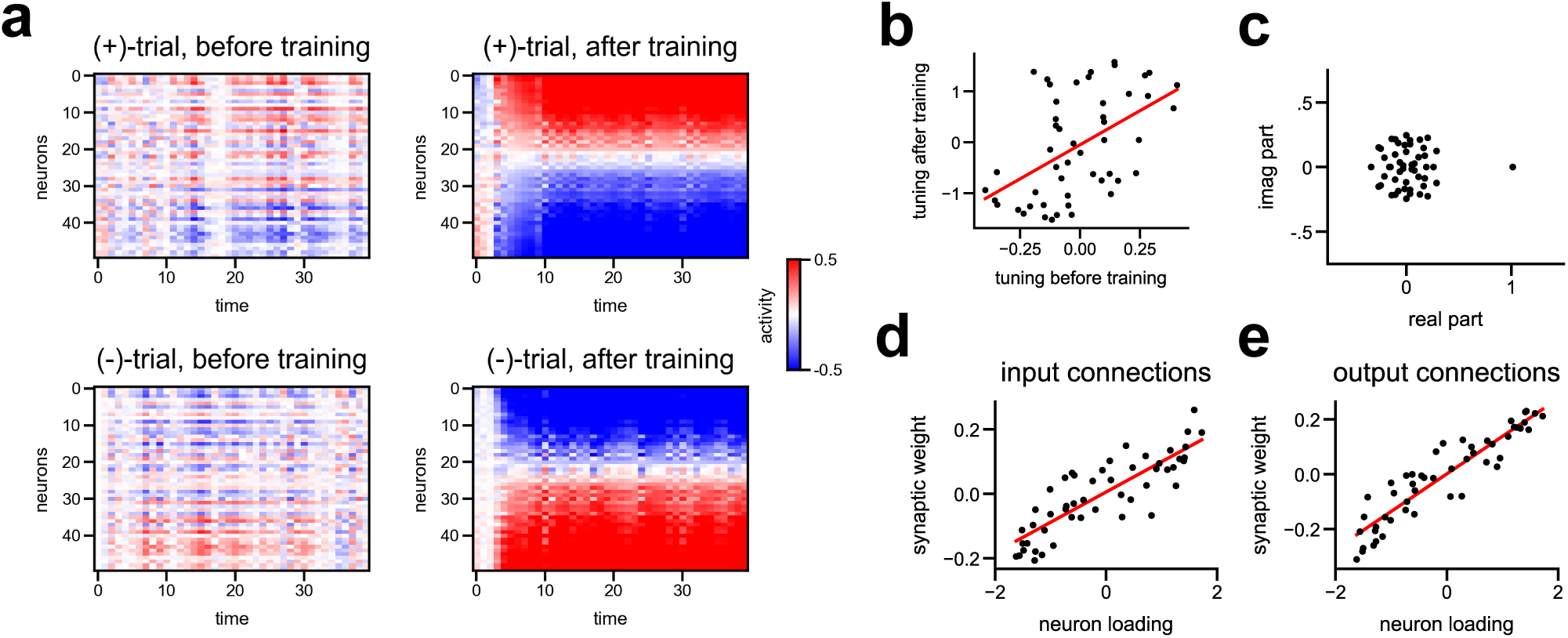
Cell tuning and synaptic connectivity properties in a nonlinear RNN trained on a stimulus discrimination task. **(a)** Activity of all cells on (+)-trials and (-)-trials before and after training. Cells were sorted by the low-dimensional neuron factor, 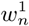, as in fig. 3e. **(b)** Cell tuning quantified as peak activity on (+)-trials minus peak activity on (-)-trials before and after training (averaged over ten trials). Cells with positive tuning scores are (+)-cells, while cells with negative tuning scores are (-)-cells. The initial tuning was positively correlated with final tuning for each cell. **(c)** Eigenvalues of the synaptic connectivity matrix after training. Similar to the solution in linear networks [53], the connectivity matrix has a single eigenvalue near 1 + 0*i*; and all other eigenvalues are small in magnitude. **(d-e)** The neuron factor identified by a 1-component TCA model is positively correlated with the input-to-network synaptic weights **(d)**, and the network-to-output weights **(e)**.

**Figure 4, Supplement 1.**
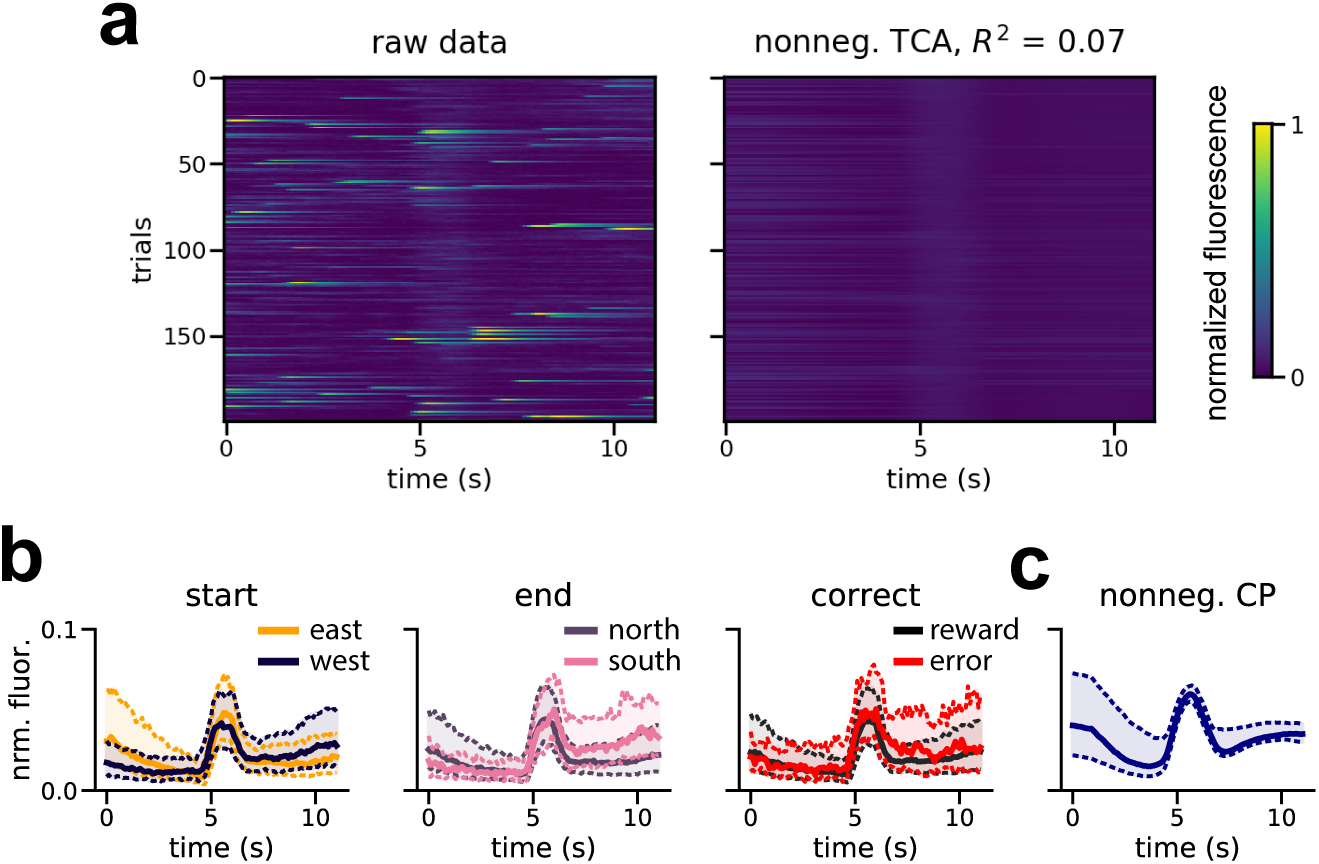
An example cell with low *R*^2^. **(a)** Raster heatmaps showing the cell’s fluorescence over the 200 most active trials (left), and the estimate of a 15-component nonnegative TCA model on these trials (right). On a small subset of trials the cell is active, but at variable phases of the trial. Note that on the remaining trials, the cell was hardly active at all (not shown). **(b)** Median fluorescence traces averaged over various task variables (start location, end location, and reward delivery). The cell does not, on average, show a preference for any task variable. Dashed lines denote the first and third quartiles of the fluorescence trace. **(c)** Median estimated fluorescence of the 15-component nonnegative TCA model for this cell. The estimate is closely matched to the median firing rates shown in panel b. Dashed lines denote first and third quartiles.

**Figure 5, Supplement 1.**
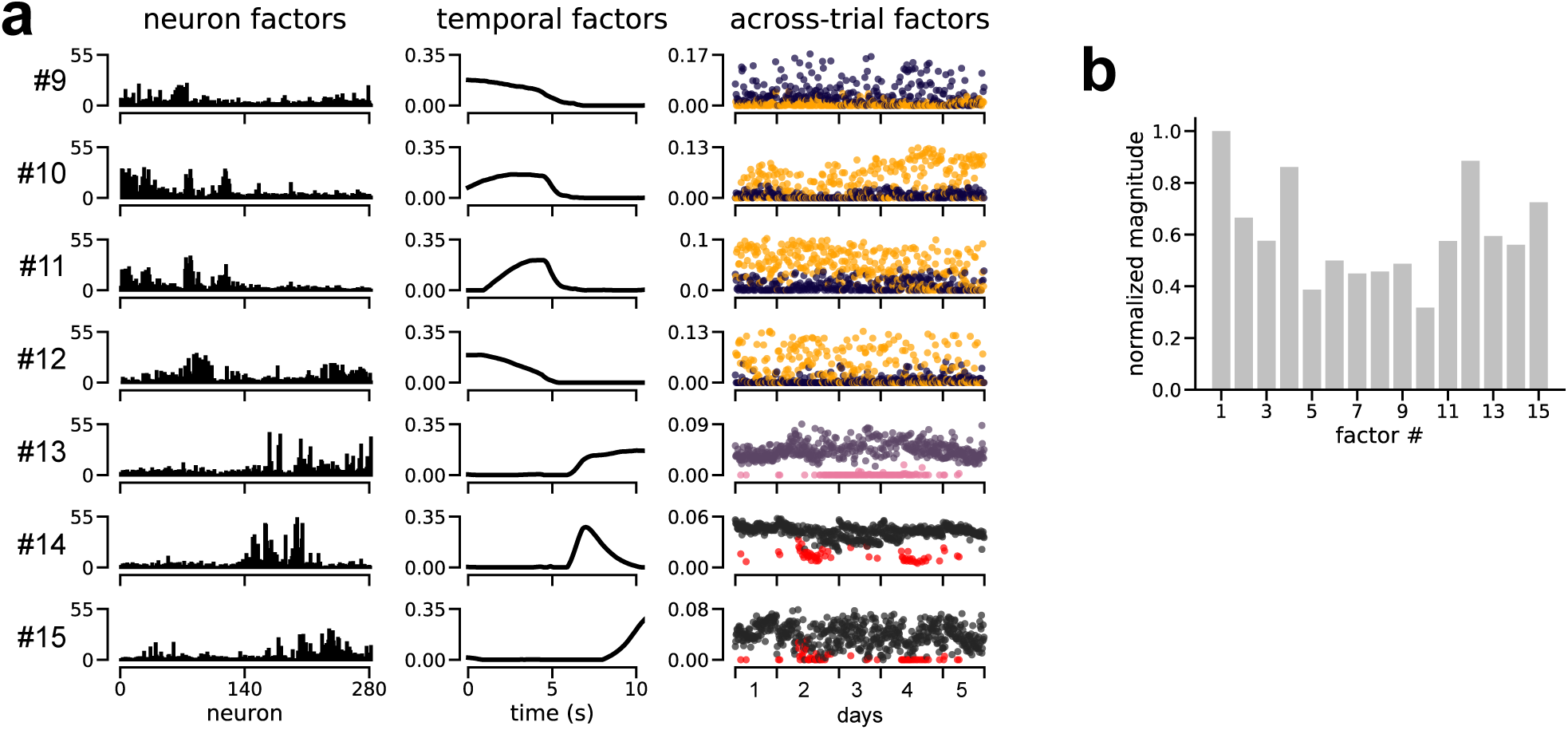
Additional detail on the decomposition of mouse prefrontal cortex dynamics. **(a)** Remaining seven TCA factors from the 15-component decomposition shown in fig. 5. **(b)** The magnitude (Euclidean length) of each factor in the decomposition, a metric analogous to the variance explained by each component (see *Methods*, section 4.5.1).

**Figure 5, Supplement 2.**
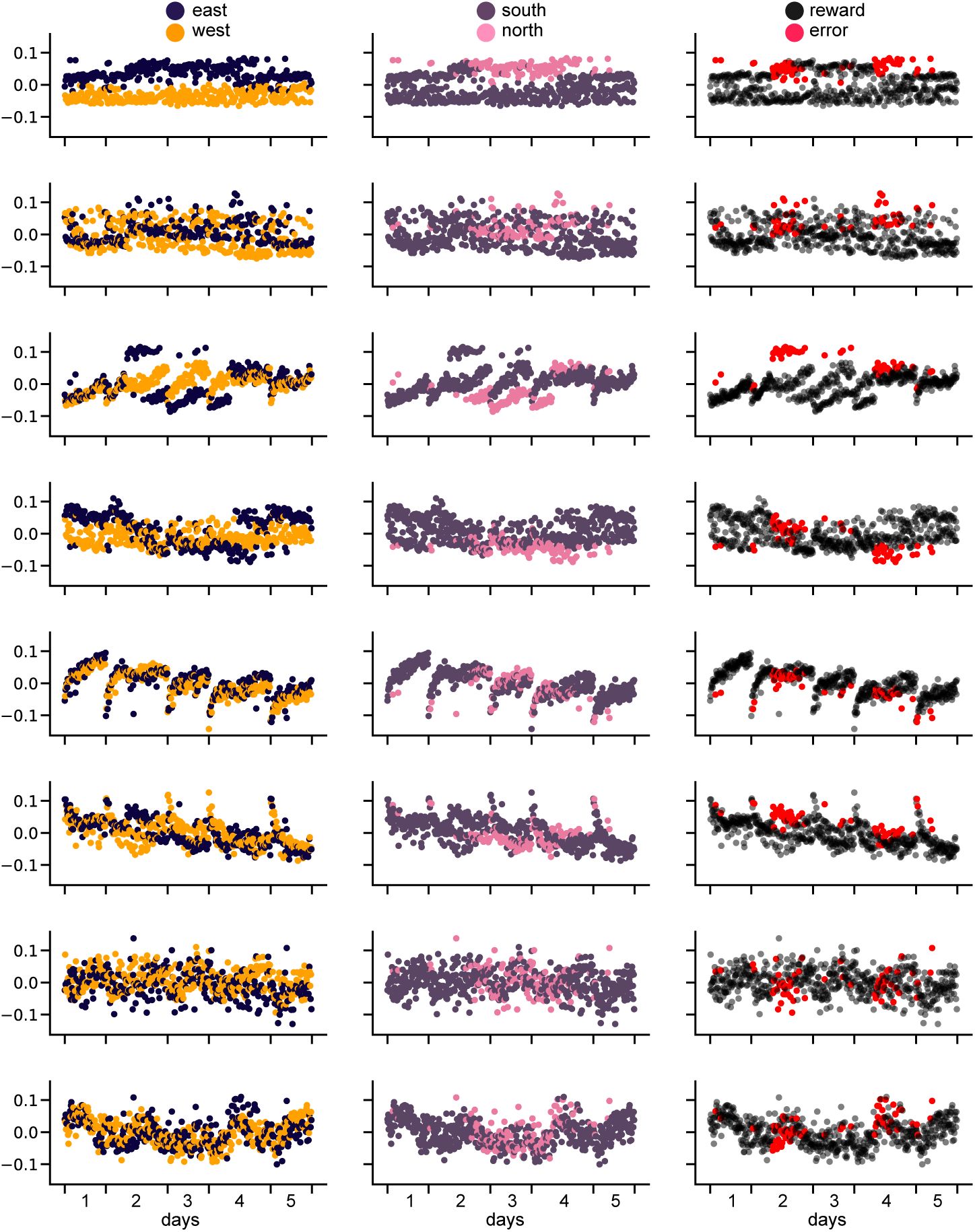
PCA components in trial-space do not cleanly encode individual task variables, in line with previous observations [34]. Each row shows a principal component, ordered by variance explained. Each column shows a different coloring of that principal component by a different task variable. With few exceptions (notably the top component), any single coloring does not yield a simple interpretation of the component.

**Figure 6, Supplement 1.**
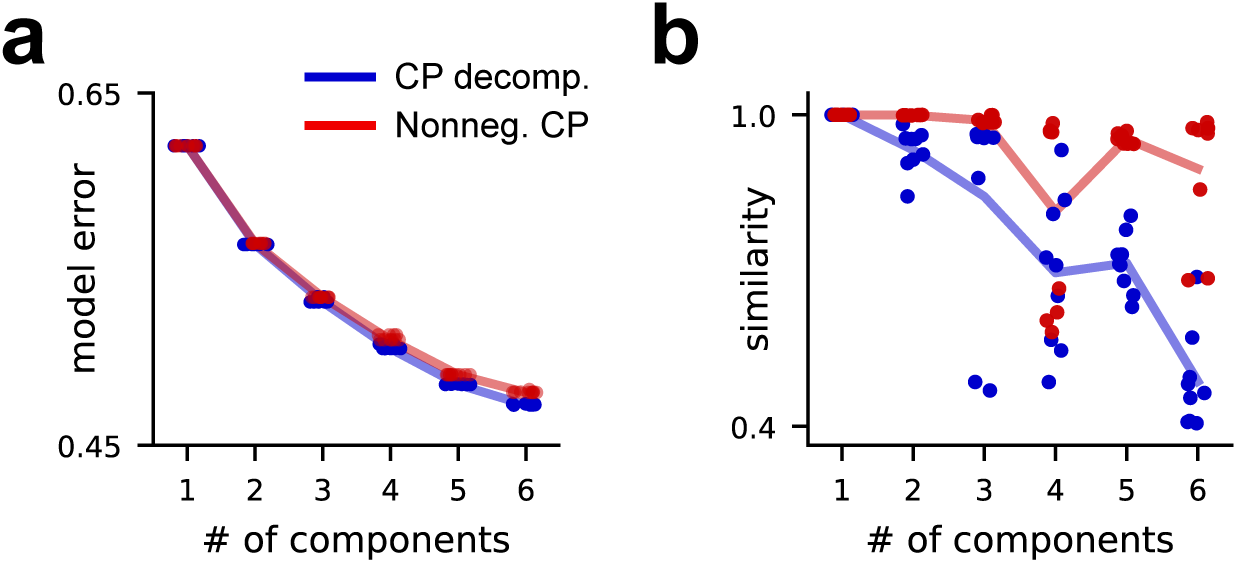
Diagnostic plots for TCA models fit to 45° reaches in the primate BMI dataset. **(a)** Scree plot for unconstrained (blue) and nonnegative (red) TCA. As elsewhere in this manuscript, each dot denotes a model fit from different initial parameters, demonstrating that neither model got caught in appreciably sub-optimal local minima during optimization. Nonnegative decomposition provided similar explanatory power to unconstrained decompositions. **(a)** Similarity plot for unconstrained (blue) and nonnegative (red) CP decompositions. As elsewhere in this manuscript, each dot denotes the similarity score between a model and the best-fit model with the same number of components. Nonnegative decomposition had larger similarity scores, suggesting that the latent factors were more reliably identified and less sensitive to initialization.

